# Gene co-expression network analysis in human spinal cord highlights mechanisms underlying amyotrophic lateral sclerosis susceptibility

**DOI:** 10.1101/2020.08.16.253377

**Authors:** Jerry C. Wang, Gokul Ramaswami, Daniel H. Geschwind

## Abstract

Amyotrophic lateral sclerosis (ALS) is a neurodegenerative disease defined by motor neuron (MN) loss. Multiple genetic risk factors have been identified, implicating RNA and protein metabolism and intracellular transport, among other biological mechanisms. To achieve a systems-level understanding of the mechanisms governing ALS pathophysiology, we built gene co-expression networks using RNA-sequencing data from control human spinal cord samples, identifying 13 gene co-expression modules, each of which represents a distinct biological process or cell type. Analysis of four RNA-seq datasets from a range of ALS disease-associated contexts reveal dysregulation in numerous modules related to ribosomal function, wound response, and leukocyte activation, implicating astrocytes, oligodendrocytes, endothelia, and microglia in ALS pathophysiology. To identify potentially causal processes, we partitioned heritability across the genome, finding that ALS common genetic risk is enriched within two specific modules, SC.M4, representing genes related to RNA processing and gene regulation, and SC.M2, representing genes related to intracellular transport and autophagy and enriched in oligodendrocyte markers. Top hub genes of this module include ALS-implicated risk genes such as KPNA3, TMED2, and NCOA4, the latter of which regulates ferritin autophagy, implicating this process in ALS pathophysiology. These unbiased, genome-wide analyses confirm the utility of a systems approach to understanding the causes and drivers of ALS.

## Introduction

Amyotrophic lateral sclerosis (ALS) is an adult-onset neurodegenerative disease defined by motor neuron (MN) loss and results in paralysis and eventually death 3-5 years following initial disease presentation^1^. In 5-10% of cases, patients exhibit a Mendelian pattern of inheritance and the disease segregates within multigenerational affected families (familial ALS)^2^. However, in 90% of cases, the underlying cause of ALS is unknown and disease occurrence is classified as sporadic (sporadic ALS)^2^. Most genetic risk in ALS is polygenic, indicating a complex genetic architecture^3^. Given this complex genetic architecture with contributions of rare and common alleles, we took a systems level approach to understand the gene expression signatures associated with ALS pathophysiology.

Weighted gene co-expression network analysis (WGCNA)^4^ has been a powerful method for organizing and interpreting wide-ranging transcriptional alterations in neurodegenerative diseases. In Alzheimer’s disease (AD), transcriptomic and proteomic studies have revealed a pattern of dysregulation implicating immune- and microglia-specific processes^5,6^. More broadly across tauopathies including AD, frontotemporal dementia (FTD), progressive supranuclear palsy (PSP), molecular network analyses have identified gene modules and microRNAs mediating neurodegeneration, highlighting its utility in prioritizing molecular targets for therapeutic interventions^7–10^. In ALS, previous studies have studied the transcriptional landscape in sporadic and C9orf72 (familial) ALS post-mortem brain tissues, finding a pattern of dysregulation in gene expression, alternative splicing, and alternative polyadenylation that disrupted RNA processing, neuronal functioning, and cellular trafficking^11^. However, the generalizability of these findings is limited due to relatively small sample sizes (n=26). Numerous studies have previously examined the transcriptional landscape of the spinal cord, which hosts axons from the upper motor neurons (UMN) and somae of the lower motor neuron (LMN) and is the location of LMN degeneration in ALS^12–17^. However, a rigorous interrogation of gene co-expression networks in the spinal cord has not yet been fully explored.

To investigate gene expression signatures associated with ALS in the spinal cord, we generated gene co-expression networks using RNA-sequencing of 62 neurotypical human cervical spinal cord samples from the GTEx consortium^18^. We characterized these co-expression modules through gene ontology (GO) enrichment analysis, which reveals that the modules represent a diverse range of biological processes. To identify modules enriched with ALS genetic risk factors, we queried enrichment of rare, large effect risk mutations previously described in the literature, as well as common genetic variation^19,20^. The second approach is genome-wide and therefore unbiased, which focuses on the enrichment of small effect common risk variants identified from a Genome Wide Association Study (GWAS) of ALS^21^. By partitioning SNP based heritability and performing enrichment testing for genes with rare mutations, we find significant overlap of genes harboring rare and common variation in SC.M4, which represents genes involved in RNA processing and epigenetic regulation. We also find enrichment of ALS common genetic risk variants in SC.M2, which represents genes involved in intracellular transport, protein modification, and cellular catabolic processes. We extend these results by demonstrating the dysregulation of these modules in data from multiple published human ALS patient and animal model tissues^12,13,22,23^. We identify modules related to ribosomal function, wound response, and leukocyte activation that are dysregulated across this diverse set of ALS-related datasets. The set of genes underlying these modules provides a rich set of targets for exploring new mechanisms for therapeutic development.

## Results

### Co-expression network analysis as a strategic framework for elucidating ALS susceptibility

We reasoned that because ALS preferentially affects spinal cord and upper motor neurons, co-expression networks from spinal cords of control subjects would provide an unbiased context for understanding potential convergence of ALS risk genes. We used the Genotype-Tissue Expression (GTEx) Project^18^ control cervical spinal cord samples to generate a co-expression network as a starting point for a genome wide analysis to identify specific pathways or cell types affected by ALS genetic risk. First, we identified gene co-expression modules and assigned each module a biological, as well as cell-type identity^24,25^. Next, we leveraged multiple datasets identifying ALS genetic risk factors^19–21,26–28^ to comprehensively assess which pathways and cell-types are susceptible to ALS genetic risk. Finally, we compared these predictions with human ALS and animal model datasets^12,13,22,23^ to develop a mechanistic understanding of ALS etiology and progression (**Fig. 1**).

**Figure 1.**
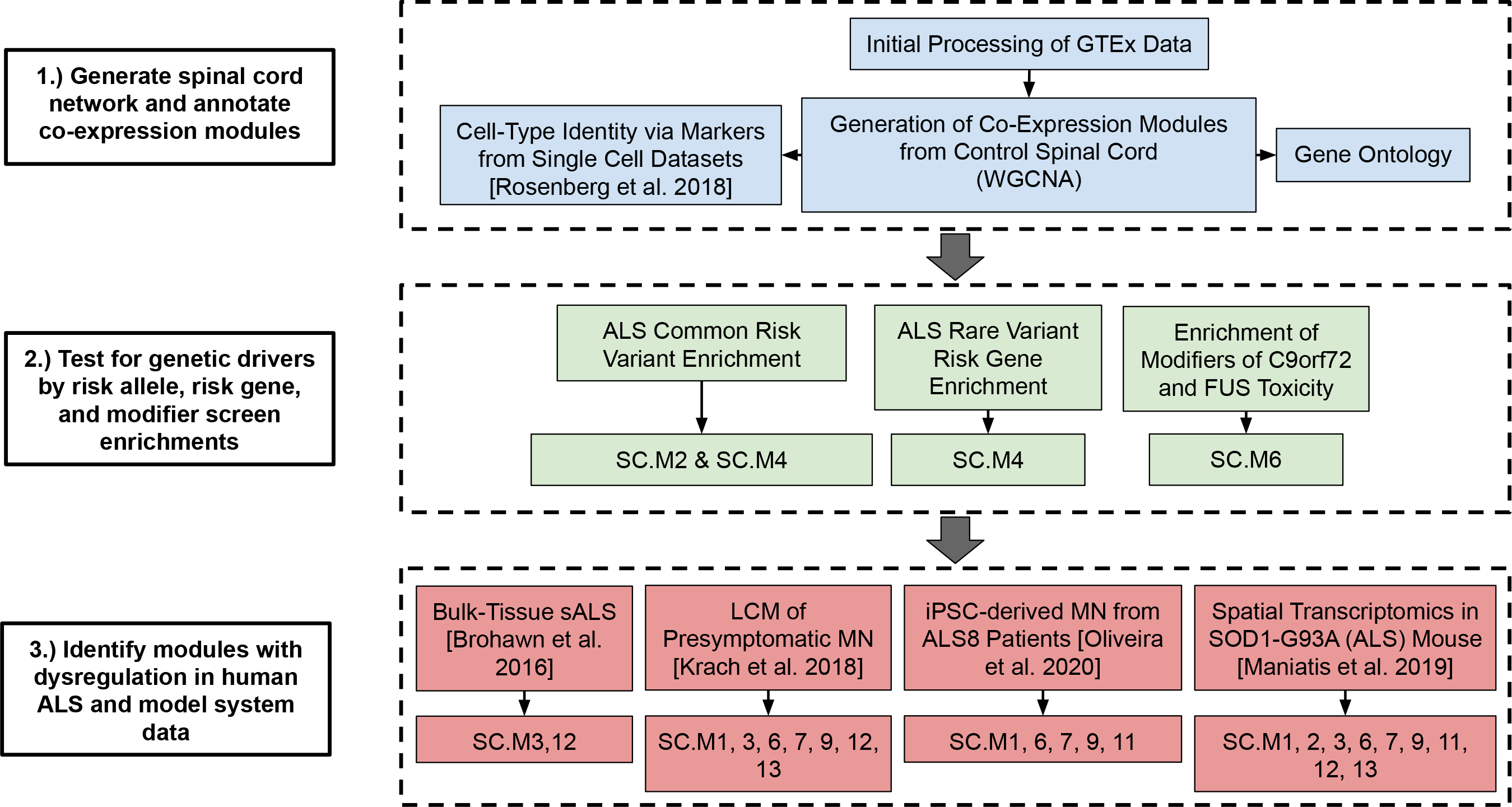
Overview of spinal cord co-expression network analysis. Flow diagram of analyses in this study. A co-expression network was generated using the GTEx cervical spinal cord expression data and Weighted Gene Co-Expression Network Analysis (WGCNA). Modules were annotated by their associated biological processes, hub genes, and cell-type identities. Modules were then assessed for their enrichment of ALS genetic risk factors and analyzed for differential expression in RNA-seq data from human ALS tissues and model system studies^12,13,22,23^.

### Generation of human spinal cord gene co-expression network

We utilized RNA-sequencing data from the GTEx consortium to generate a co-expression network of human spinal cord (**Methods**). Co-expression networks are highly sensitive to biological and technical confounders^25^, which we controlled for by calculating the correlation of biological and technical covariates with the top expression principal components (PCs) (**Methods, Supplementary Fig. S1a-c**). We identified 11 technical and biological covariates: seqPC1-5, RNA integrity number (RIN), Hardy’s death classification scale, age, sampling center, ethnicity, and ischemic time as significant drivers of expression (**Supplementary Fig. S1a**). We regressed out the effect of these covariates using a linear model (**Methods**) and verified that the regressed expression dataset was no longer correlated with technical and biological covariates (**Supplementary Fig. S1b**).

We applied WGCNA to construct a co-expression network (**Fig. 2a, Supplementary Fig. S2; Supplementary Data 1; Methods**) identifying 13 modules. To verify that the modules were not driven by confounding covariates, we confirmed that the module eigengenes were not correlated with the major technical and biological covariates (**Supplementary Fig. S1d**). We also found that randomly sampled genes from within each module were significantly more inter-correlated when compared to a background set, verifying the co-expression of genes within each module (**Supplementary Fig. S1e**). To biologically annotate our co-expression modules, we performed gene ontology (GO) term enrichment and identified the top hubs (**Fig. 2c-g, 3c-d, Supplementary Fig. S3**). GO analysis revealed that co-expression modules represented coherent biological processes that we categorized into five distinct groupings (**Fig. 2b**), encompassing morphogenesis and development, epigenetics and gene regulation, neuronal signaling, immune activation, and intracellular transport, metabolism, and sensing functions. Most modules were significantly enriched for markers of specific cell types, suggesting that they represent gene co-expression within major cell classes, including SC.M1 (astrocytes), SC.M2 (oligodendrocytes), SC.M3 (endothelial cells), SC.M9 and SC.M11 (neurons), SC.M10 (ependymal), and SC.M12 and SC.M13 (microglia) (**Supplementary Fig. S4**). These groupings, both with regards to cell type and molecular pathways, represent diverse biological functions that play key roles in human spinal cord biology.

**Figure 2.**
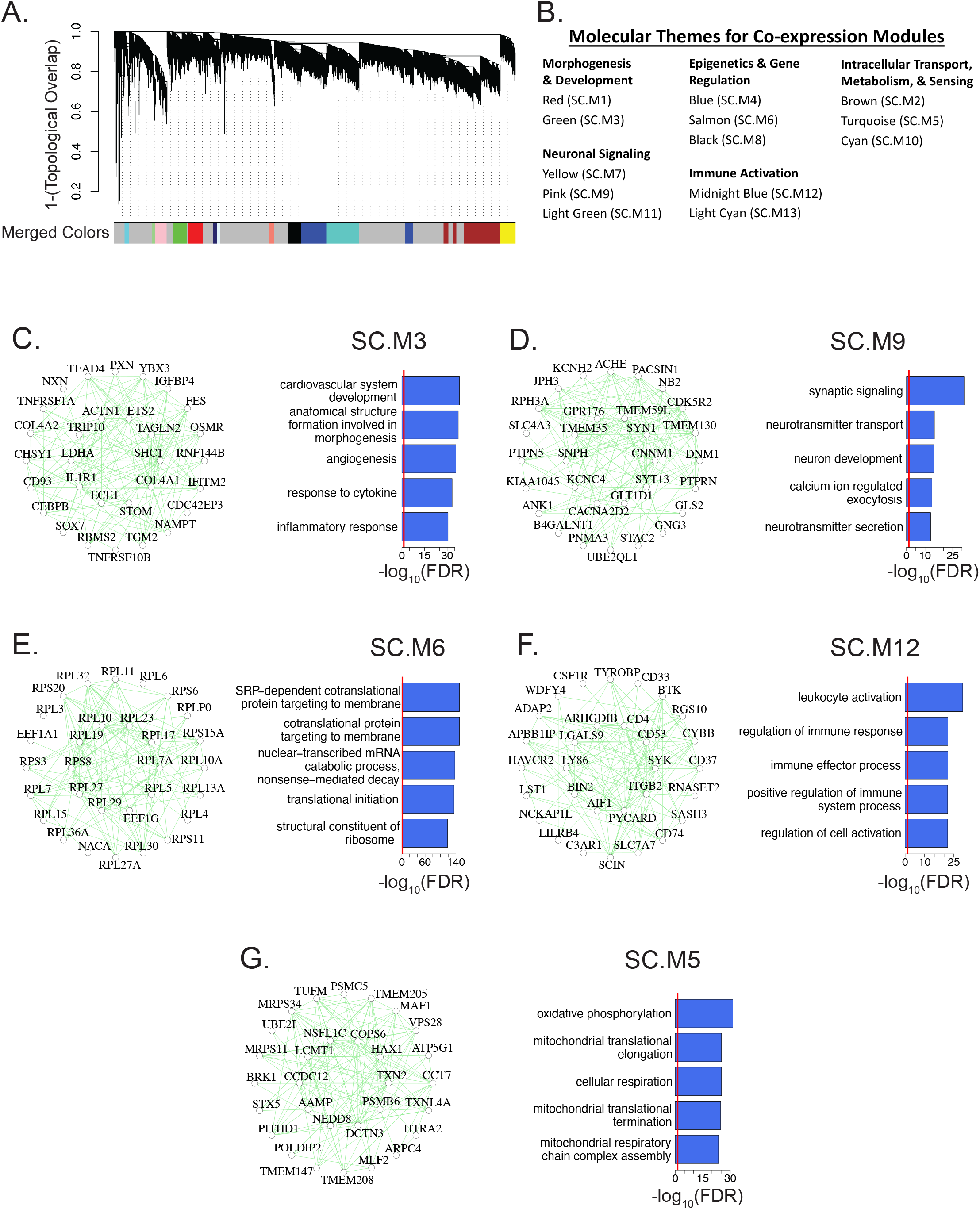
Construction of co-expression network in control cervical spinal cord. (A) Dendrogram of the topological overlap matrix for gene co-expression dissimilarity. Co-expression module assignments are shown in the “Merged Colors” track (See Supplementary Fig. S2 for optimization of tree-cutting parameters). Genes that failed to be clustered into a co-expression module were grouped within the grey module. (B) Thirteen co-expression modules were categorized into five broad molecular themes based on Gene Ontology (GO) term enrichment and assigned a unique module identifier (SC.M1-13). (C, D, E, F, G) Top 30 hub genes and 300 connections for SC.M3, SC.M9, SC.M6, SC.M12, and SC.M5 respectively (left panels). Top 5 enriched GO terms for each of the respective modules (right panels). Red lines mark an FDR corrected p-value threshold of 0.05.

### Enrichment of ALS genetic risk in intracellular transport/autophagy module SC.M2 and RNA processing/gene regulation module SC.M4

To address how these modules fit within the context of known ALS susceptibility factors, we assessed whether a literature-curated set of well-known, large effect size, familial ALS risk genes^19,20^ were enriched within our co-expression modules (**Fig. 3a**). We found a nominally significant enrichment with SC.M4 (OR = 2.52, P = 0.0124). However, the FDR adjusted p-value was not significant (FDR= 0.174), likely due to the small number of familial ALS risk genes identified to date. SC.M4 was highly enriched with GO terms related to RNA processing and epigenetic regulation (**Fig. 3c**), consistent with the known role of several ALS risk genes in RNA metabolism and function. To prioritize interacting partners to ALS risk genes found in SC.M4, we leveraged a direct protein-protein interaction (PPI) network using the top 300 hub genes of SC.M4, finding a significant network that includes ALS risk genes such as TARDBP and EWSR1 (**Fig. 3e, Methods**, p = 0.001).

**Figure 3.**
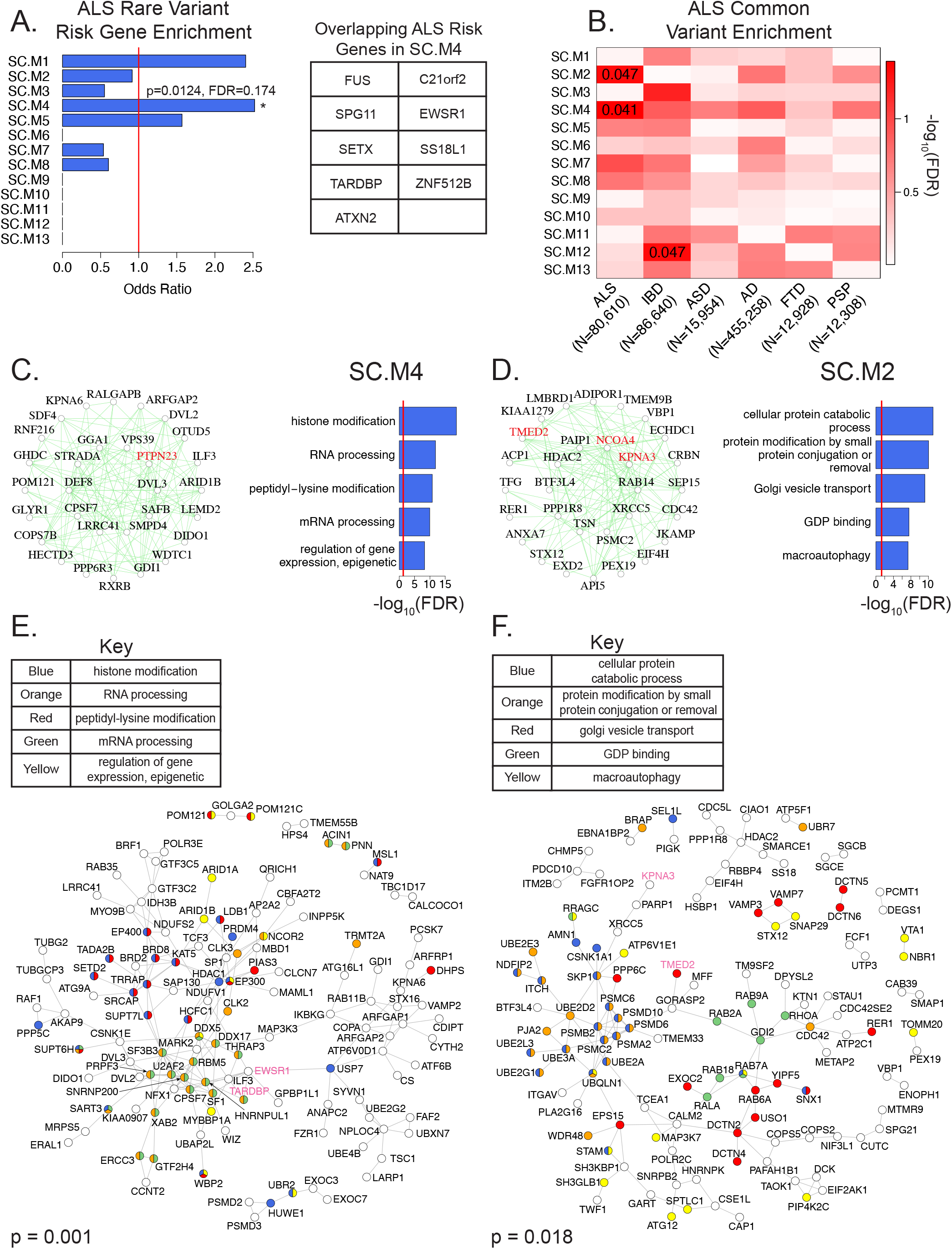
Enrichment of ALS genetic risk factors in intracellular transport/autophagy module SC.M2 and RNA processing/gene regulation module SC.M4. (A) Enrichment of literature curated large-effect ALS risk genes^19,20^ within co-expression modules (left panel). Red line marks a baseline odds ratio of 1. SC.M4 is marked with an asterisk as having a nominal enrichment p-value < 0.05. Literature-curated large-effect ALS risk genes within the SC.M4 module (right panel). (B) Partitioned heritability enrichments of common risk variants for ALS^21^, IBD^30^, ASD^32,33^, AD^34^, FTD^35^, and PSP^36^ within co-expression modules. Text values of enrichments with FDR corrected p-value < 0.05 are shown. (C, D) Top 30 hub genes and 300 connections for SC.M4 and SC.M2 respectively (left panels). Highlighted in red are genes that have been previously implicated in ALS pathophysiology^27,43–45^. Top 5 enriched GO terms for each of the respective modules (right panels). Red lines mark an FDR corrected p-value threshold of 0.05. (E, F) Direct protein-protein interaction (PPI) network of the top 300 hub genes from modules SC.M4 and SC.M2 respectively. Gene vertices are color coded according to their Gene Ontology terms as shown^63^. P-value of the PPI network is derived from 1,000 permutations. Highlighted in pink are genes that have been previously implicated in ALS pathophysiology^27,43–45^.

The small number (∼50) of large effect size risk genes identified to date in ALS makes the detection of strong enrichments difficult. To complement the rare risk gene enrichment analyses, we performed an unbiased genome-wide analysis, partitioning the common SNP-based heritability across the co-expression modules using stratified LD score-regression^29^ based on summary statistics from the most recent GWAS of ALS (ALS n= 20,806; CTL n = 59,804)^21^ (**Fig. 3b**). We found significant enrichment of ALS common risk variants in the SC.M2 (FDR = 0.047) and SC.M4 modules (FDR = 0.041). SC.M2 was enriched with GO terms related to intracellular transport, protein modification, and cellular catabolic processes (**Fig. 3d**). Additionally, SC.M2 was significantly enriched with oligodendrocyte markers (**Supplementary Fig. S4**, Oligodendrocytes_mature: OR = 5.3, FDR < 10e-5; Oligodendrocytes_myelinating: OR = 2.9, FDR < 0.05), while SC.M4 showed no specific cell type enrichment. Generation of a direct protein-protein interaction (PPI) network using the top 300 hub genes of SC.M2 reveals a significant network that highlights SC.M2 hub genes that interact with known ALS risk genes such as TMED2 and KPNA3^27^ (**Fig. 3f, Methods**, p = 0.018).

As a comparison, we tested for the enrichment of common risk variants from Inflammatory Bowel Disorder (IBD)^30^ (N=86,640) across modules (**Fig. 3b**). We found a significant enrichment of IBD common risk variants (FDR = 0.047) in module, SC.12, which is a microglial/immune module, consistent with the known disease biology of IBD^31^. As a negative control, we performed enrichments of common risk variants from Autism Spectrum Disorder (ASD)^32,33^ (N=15,954) and found no enrichment in any module. Further, when we compared with Alzheimer’s Disease (AD)^34^ (N=455,258), Frontotemporal Dementia (FTD)^35^ (N=12,928), and Progressive Supranuclear Palsy (PSP)^36^ (N=12,308) (**Fig. 3b**), we also found no significant enrichments for these diseases’ common risk variants, demonstrating the specificity of the ALS GWAS enrichment for modules SC.M2 and SC.M4. Overall, these genetic enrichment analyses implicate dysfunction in RNA processing and intracellular transport, including in oligodendrocytes, as putative drivers of ALS pathophysiology in the human spinal cord.

### Enrichment of ALS genetic modifiers in ribosomal-associated module SC.M6

As an orthogonal approach to assess which genes and pathways are mediators of ALS pathophysiology, we leveraged genetic screen datasets from three studies of modifiers of C9orf72 and FUS toxicity in yeast^26–28^. We assessed our human spinal cord modules for enrichment with modifiers of C9orf72 and FUS toxicity identified from these screens. We found significant enrichment of C9orf72 and FUS toxicity suppressors in SC.M6 (**Fig. 4a**), a module representing genes involved in ribosomal function (**Fig. 2e**), but without any cell type specific enrichment (**Supplementary Fig. S4**). This finding is consistent with previous observations that modifiers of Glycine-Arginine_100_ (GR_100_) DPR and FUS toxicity are enriched with gene ontology terms related to ribosomal biogenesis^26,28^.

**Figure 4.**
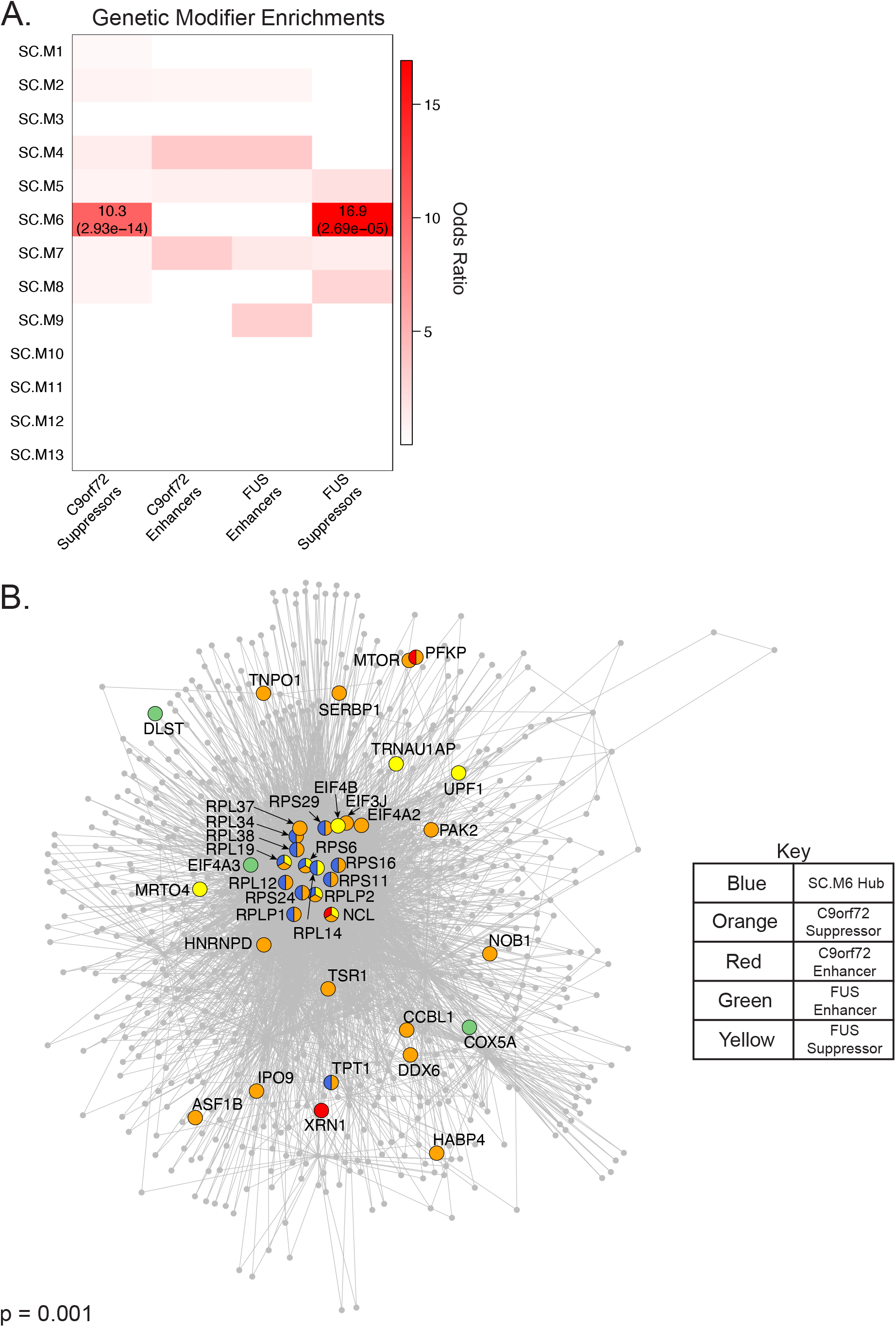
Modifiers of C9orf72 and FUS toxicity converge on top hubs of ribosomal-associated module SC.M6. (A) Enrichment of genetic modifiers of C9orf72 and FUS toxicity^26–28^ within co-expression modules. Odds ratios of enrichments with FDR corrected p-value < 0.05 are shown. FDR corrected p-values themselves are shown within parentheses. (B) Direct protein-protein interaction (PPI) network of the top 100 hub genes from module SC.M6. Gene vertices are color coded as shown. P-value of the PPI network is derived from 1,000 permutations.

We reasoned that SC.M6 could be leveraged to prioritize highly interconnected hub genes within the module that may warrant additional scrutiny as points of convergence for C9orf72 and FUS pathology. To address this, we first identified the interacting partners for the top 100 SC.M6 hub genes by generating an indirect protein-protein interaction (PPI) network (**Methods**). We identified 30 modifiers of C9orf72 and FUS toxicity that were present in this indirect PPI network, either as hubs or as interacting partners (**Fig. 4b**)^26–28^. These modifiers include ribosomal proteins (RPL and RPS-related genes), intracellular transporters of ribosomal components (TNPO1), ribosomal assembly proteins (NOB1), and RNA-binding proteins (HABP4) (**Fig. 4b**), indicating specific aspects of ribosomal dysfunction shared between C9orf72 and FUS toxicity. Indeed, previous studies^26^ have suggested that the overlap in modifiers of C9orf72 and FUS toxicity could be due to the sequence similarity between the arginine/glycine/glycine (RGG) domain in FUS and the GR_100_ sequence expressed by C9orf72^37,38^.

### Dysregulation of immune-related modules in postmortem ALS spinal cord

Until this point, we had assessed co-expression modules for their relevance to ALS based on enrichment of genetic risk factors. To assess whether the co-expression modules identified using the GTEx dataset were directly dysregulated in ALS patients, we processed bulk RNA-sequencing data from a previous study of postmortem control and ALS cervical spinal cord^13^. We corrected the data for potential confounders such as age, ethnicity, sex, and sequencing variability and assessed module preservation (**Supplementary Fig. S5a-b, Methods**). We identified 10 of the 13 modules generated from GTEx as preserved in the postmortem control and ALS cervical spinal cord (Z_summary_ > 2) with four (SC.M2, SC.M4, SC.M5, and SC.M8) strongly preserved (Z_summary_ > 10). The remaining three modules (SC.M9, SC.M11, and SC.M10) were not preserved (Z_summary_ < 2) (**Supplementary Fig. S5c**).

We adopted two approaches to assess which modules were dysregulated in ALS in the human spinal cord. The first assessed enrichments of differentially expressed genes (DEGs) between postmortem control and ALS cervical spinal cord samples^13^ (**Methods**). We found a significant enrichment of DEGs upregulated in cervical spinal cord from ALS patients in SC.M12 (FDR = 5.92e-39) and of downregulated DEGs in SC.M3 (FDR = 0.00152) (**Fig. 5a**). SC.12 is highly enriched with microglial markers and GO terms related to immune response (**Fig. 2f, Supplementary Fig. S4a**), whereas SC.M3 is highly enriched with endothelial markers and GO terms related to inflammation and morphogenesis (**Fig. 2c, Supplementary S4a**).

**Figure 5.**
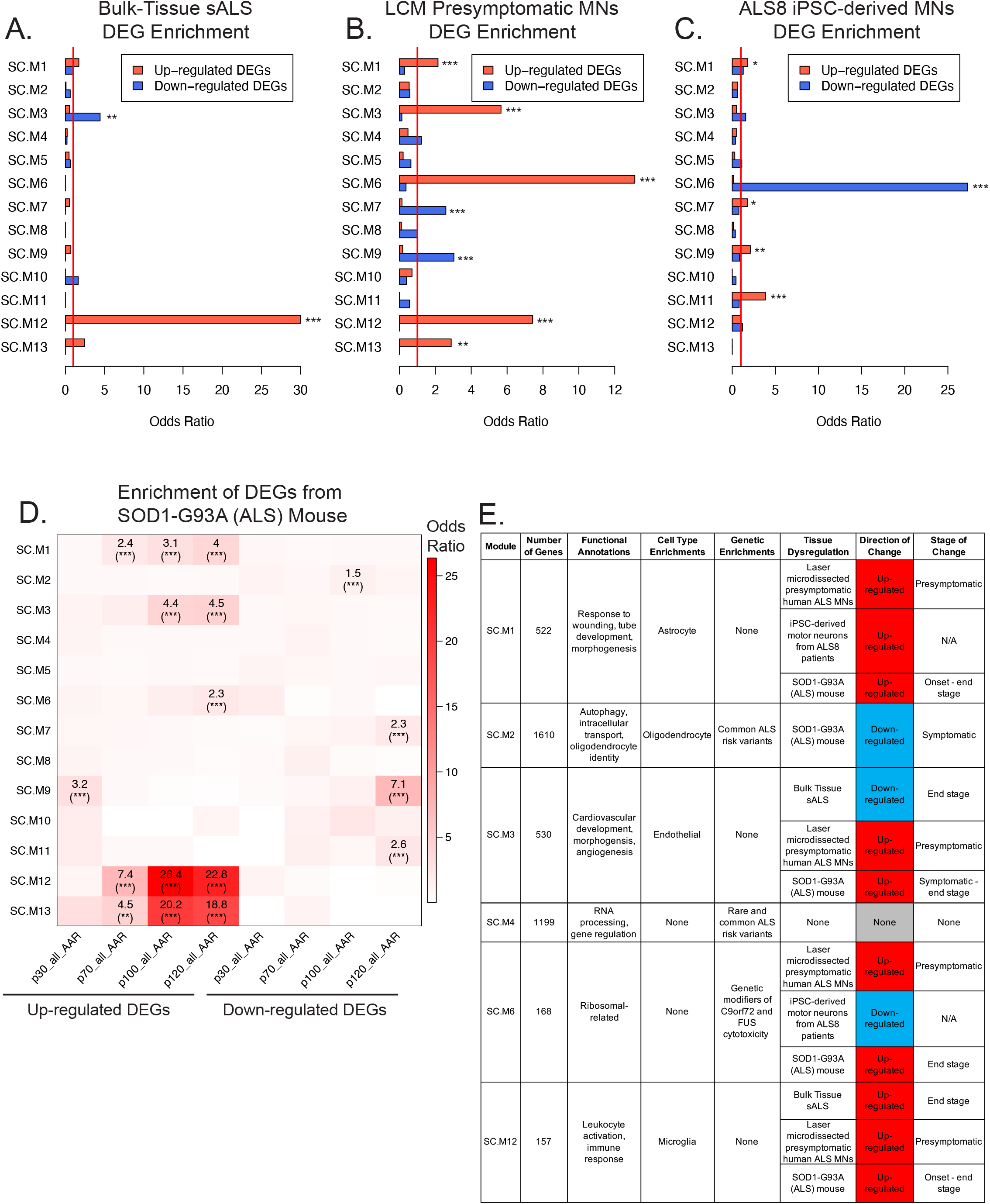
Validation of co-expression module dynamics in human ALS and model system datasets. (A) Enrichments of 112 up-regulated and 70 down-regulated DEGs from bulk-tissue RNA-sequencing of postmortem ALS cervical spinal cord^13^ within co-expression modules. Red line marks a baseline odds ratio of 1. * (FDR corrected p-value < 0.05). ** (FDR corrected p-value < 0.01). *** (FDR corrected p-value < 0.001). (B) Same as Fig. 5a except for using 1290 up-regulated and 310 down-regulated DEGs from RNA-sequencing of laser capture microdissections of motor neurons primed for degeneration ^22^. (C) Same as Fig. 5a except for using 364 up-regulated and 289 down-regulated DEGs from RNA-sequencing of iPSC-derived motor neurons from ALS8 patients^23^. (D) Enrichment of DEGs within each disease stage of the SOD1-G93A mouse^12^ within co-expression modules. p30, p70, p100, and p120 represent postnatal age in days at time of spatial transcriptomic sequencing. Odds ratios of enrichments with FDR corrected p-value < 0.05 are shown. * (FDR corrected p-value < 0.05). ** (FDR corrected p-value < 0.01). *** (FDR corrected p-value < 0.001). (E) Summary of the key findings from this study regarding modules relevant to ALS pathophysiology.

For our second approach, we assessed if the spinal cord modules were themselves differentially expressed between ALS and control samples by looking at the relationship between the eigengenes (first principle component; **Methods**) of each module and disease status (**Supplementary Fig. S5d**). We found that the SC.M1 eigengene was significantly upregulated in ALS samples. SC.M1 was enriched with astrocyte markers and GO terms related to wound response (**Supplementary Fig. S3a, S4a**). Overall, our analyses identify a marked upregulation of neuro-immune cells in the human post-mortem spinal cord, including astrocytes and microglia, similar to previous reports^13,22^. However, we note that genes differentially expressed in postmortem ALS spinal cord are not enriched in causal genetic risk, suggesting that these changes are not causal drivers.

### Dysregulation of immune and neuronal related modules in dissected human motor neurons primed for degeneration

A major disadvantage of utilizing postmortem human tissues is that the molecular pathology is indicative of late-stage disease. In particular, immune hyperactivation and infiltration is consistently seen in neurodegenerative disorders including ALS. However, it is unclear if these processes are disease causing or merely a reaction to neurodegenerative cell death^39^. To identify early-stage differences in human spinal cord potentially representative of causal factors, we leveraged a previously published dataset of laser capture micro-dissected motor neurons deemed to be primed for degeneration^22^. In this study, the authors isolated motor neurons from ALS patients’ postmortem spinal cord, but in regions that did not exhibit any signs of degeneration at the time of death. We processed and corrected the data for potential confounders such as age, gender, postmortem interval, RNA integrity, and sequencing variability^22^ (**Supplementary Fig. S6a-b**). We identified 9 of the 13 modules generated from GTEx as preserved in the micro-dissected motor neuron dataset (Z_summary_ > 2) with two (SC.M3 and SC.M6) strongly preserved (Z_summary_ > 10). The remaining four modules (SC.M2, SC.M4, SC.M8, and SC.M10) were not preserved (Z_summary_ < 2) (**Supplementary Fig. S6c**).

We assessed the enrichment of DEGs between ALS and control micro-dissected motor neurons in our modules. We found enrichment of up-regulated DEGs within SC.M1, SC.M3, SC.M6, SC. M12, and SC.M13, modules related to astrocyte, endothelial, and microglial functions (**Supplementary Fig. S4a**). We found enrichment of down-regulated DEGs within SC.M7 and SC.M9 (**Fig. 5b**), two modules related to synaptic signaling (**Fig. 2d, Supplementary Fig. S3b**), possibly marking degeneration at early stages of ALS. We also assessed the relationship between the module eigengene and disease status, finding significant up-regulation of SC.M1, SC.M3, SC.M5, SC.M6, SC.12, and SC.M13 in the ALS samples (**Supplementary Fig. S6d**). Despite performing laser-capture micro-dissection of motor neurons, our results indicate that the predominant signal from this dataset highlights the infiltration of neuro-immune cells, including astrocytes and microglia in the progression of ALS pathophysiology.

### Dysregulation of a ribosome-related module in iPSC-derived motor neurons from ALS8 patients

To gain further insights into processes dysregulated in motor neurons and avoid the pro-immune phenotypes observed in previous postmortem studies^13,22^, we leveraged a RNA-seq dataset generated from iPSC-derived motor neurons from ALS8 patients^23^. In this study, iPSC-derived motor neurons were differentiated from five patients carrying pathogenic variants in the VAPB gene and three unaffected control individuals. We processed and corrected this dataset for potential confounders such as age, sex, sampled tissue, and sequencing variability (**Supplementary Fig. S7a-b**). We identified 10 of the 13 modules generated from GTEx as preserved in the iPSC-derived motor neuron dataset (Z_summary_ > 2) with two (SC.M7 and SC.M9) strongly preserved (Z_summary_ > 10). The remaining three modules (SC.M5, SC.M12, and SC.M13) were not preserved (Z_summary_ < 2) (**Supplementary Fig. S7c**).

We assessed the enrichment of DEGs between iPSC-derived MNs from severe ALS versus control individuals in our modules. We found modest enrichment of up-regulated DEGs within SC.M1, SC.M7, SC.M9, and SC.M11 (OR = 1.77, 1.76, 2.09, 3.84 respectively), modules related to astrocyte and neuronal functions (**Fig. 5c, Supplementary Fig. S4**). We found a strong and pronounced enrichment of down-regulated DEGs within SC.M6 (OR = 27.28), a ribosomal-related module (**Fig. 2e, 5c**). We also assessed the relationship of module eigengene to disease status, revealing up-regulation of SC.M2 and SC.M7 and down-regulation of SC.M1 (**Supplementary Fig. S7d**). By analyzing gene expression in iPSC-derived motor neurons from ALS patients, we identified striking downregulation of ribosomal function, represented by SC.M6, as a prominent dysregulated feature specific to MNs.

### Dysregulation of immune and neuronal-related modules in SOD1-G93A (ALS) mice

Finally, as an orthogonal approach to identify temporal differences in ALS, we leveraged a spatial transcriptomics dataset from spinal cord of SOD1-G93A mice over four time points during disease progression^12,40,41^. To assess which modules were dysregulated over the progression of disease, we performed an enrichment analysis between the human spinal cord modules and mouse DEGs stratified by time point (**Methods**). Consistent with our findings in postmortem tissues (**Fig. 5a-b, Supplementary Fig. S6e**), immune response modules (SC.M12 and SC.M13) were highly enriched with up-regulated DEGs as early as postnatal day 70, which is the time of clinical disease onset in this model (**Fig. 5c, 2f, Supplementary Fig. S3f**). We also noted a strong enrichment of down-regulated DEGs in neuronal signaling modules (SC.M7, SC.M9, and SC.M11) (**Supplementary Fig. S3b, Fig. 2d, Supplementary Fig. S3e**) by postnatal day 120 (p120 - end-stage ALS), which is consistent with motor neuron degeneration. Interestingly, we found enrichment of down-regulated DEGs in SC.M2 at p100 (symptomatic ALS), providing evidence that SC.M2 is dysregulated at symptomatic stages of ALS in this model system (**Fig. 5c, Supplementary Fig. S9**). We also noted an enrichment of up-regulated DEGs in SC.M6 at p120 (end-stage ALS), suggesting dysregulation of SC.M6 in the end-stages of ALS in this model system (**Fig. 5c, Supplementary Fig. S8**). Overall, enrichment analyses in SOD1-G93A (ALS) mice highlight immune and neuronal-related modules as key disrupted pathways in the spinal cord and implicates the dysregulation of SC.M2, a module enriched with ALS genetic risk, and SC.M6, a module enriched with ALS genetic modifiers, in later stages of disease progression (**Fig. 5e**).

## Discussion

To help elucidate the biological mechanisms underlying ALS pathophysiology in an unbiased genome-wide manner, we generated co-expression networks using RNA-sequencing of control human cervical spinal cord samples from the GTEx Consortium. We identified 13 total co-expression modules each representing distinct aspects of biology in the control spinal cord including inflammation, neurotransmission, RNA processing, and cellular metabolism, as well as major cell classes. We find that ALS genetic risk factors are enriched in SC.M4, a co-expression module representing genes involved in RNA processing and epigenetics and SC.M2, which is enriched in oligodendrocyte markers and genes involved in intracellular transport, protein modification, and cellular catabolic processes.

It is notable that genes within both SC.M2 and SC.M4 have previously been shown to cause rare Mendelian forms of ALS or have been strongly linked with risk. For example, SC.M4, which is enriched in RNA processing, harbors RNA binding proteins mutated in ALS including TDP-43 and FUS/TLS, which are abnormally aggregated and mis-localized in ALS affected neurons, leading to the loss of normal RNA binding function with incitement of neurotoxicity^42^. One of the top hubs of the SC.M4 module is PTPN23, a non-receptor-type tyrosine phosphatase that is a key regulator of the survival motor neuron (SMN) complex, which helps assemble small ribonucleoproteins particles (snRNPs) and mediates pre-mRNA processing in motor neurons^43^. Furthermore, two other top hubs in SC.M2, KPNA3 and TMED2, are nucleocytoplasmic transporters that act as modifiers of dipeptide repeat (DPR) toxicity in cases of C9orf72 mutations^27^. NCOA4, another top hub in SC.M2, is a key mediator of ferritinophagy and has been implicated in neurodegeneration^44,45^. Taken together, the presence of ALS-linked risk genes as hubs in SC.M2 and SC.M4 further underscores the importance of aberrant RNA processing^42^ and intracellular transport^46^ as processes likely driving ALS pathophysiology.

We also find enrichment of genetic modifiers of ALS in SC.M6, indicating the importance of ribosomal proteins and elongation factors (**Fig. 4B**) in mitigating ALS pathophysiology. These factors have been suggested to act by reducing translation of toxic dipeptide repeat (DPRs) expressed off of the GGGGCC hexanucleotide repeat expansion found in C9orf72^26^. We found SC.M6 to be up-regulated with disease status in the SOD1-G93A mouse and dissected human motor neurons primed for neurodegeneration (**Fig. 5b, d**), datasets with predominant glial infiltration. Interestingly, we found a striking down-regulation of SC.M6 in iPSC-derived motor neurons from ALS8 patients (**Fig. 5c**), a dataset without contaminating glial signal. This suggests that role of this co-expression module is complex and cell-type dependent in ALS.

Across three of four ALS tissue and model datasets analyzed, we noted the presence of a strong pro-immune phenotype (**Fig. 5**). While immune dysregulation is a key molecular feature of the spinal cord by mid-to late-stage ALS, it remains unclear whether this disease signature is primary or secondary^47,48^. Our analyses focused on ALS genetic risk factors, which serve as causal anchors to identify, in an unbiased manner, specific modules that implicate processes involved at the early stages of ALS pathophysiology. Furthermore, immune dysregulation is a common feature in post-mortem tissue from other neurodegenerative and neuropsychiatric disorders^8,49,50^; however, the enrichment of genetic risk within upregulated immune genes is complex. For neuropsychiatric disorders such as autism and schizophrenia, the strongest genetic enrichments are within neuronal genes^49^. In contrast, the strongest genetic risk enrichments for Alzheimer’s disease are within microglia^51^. For ALS, a recent assessment of genetic risk within different cell types implicates oligodendrocytes^51^ which is consistent with our finding of risk enrichment within the oligodendrocyte-enriched module, SC.M2 (**Fig. 3b**).

A major question still remains concerning the specificity of MN degeneration in ALS. Our results support the hypothesis that ALS is a non-cell-autonomous disease^52^, as evidenced by SC.M1, SC.M2, SC.M3, SC.12, and SC.M13 being enriched with astrocyte, oligodendrocyte, endothelia, and microglia markers (**Supplementary Fig. S4**). Oligodendrocytes have critical roles through the metabolic support of neurons and neurotransmission in the central nervous system^53^. A previous study has shown that oligodendrocytes harboring familial and sporadic, but not C9orf72 variants of ALS, induce motor neuron death via a SOD1-dependent mechanism, but can be rescued via lactate supplementation^54^. These previous observations corroborate our findings that highlight the potential deleterious effect of dysregulation in the intracellular transport and autophagy system on oligodendrocyte - MN dynamics in ALS. Furthermore, neuro-immune responses mediated by astrocytes and microglia also play a critical role in ALS pathogenesis^55,56^ and disruptions of blood spinal cord barrier and damage to the endothelial cells have been observed in electron micrographs from ALS^57^. Overall, our analyses support the notion that ALS is multifactorial disorder, implicating a number of cell-types and mitigating mechanisms, acting at different stages of disease. Invariably, understanding of cell-type specific mechanisms governing MN degeneration will involve future interrogation of the transcriptome and epigenome from ALS spinal cord at single-cell resolutions. Additionally, emerging RNA-sequencing datasets, including from the TargetALS foundation (http://www.targetals.org), contain larger sample sizes of ALS and control spinal cord that will help elucidate the cellular dynamics underlying ALS pathophysiology.

## Methods

### Initial processing of RNA-sequencing datasets

The latest version of GTEx data (version 7) was obtained from the dbGaP accession number phs000424.v7.p2 on 10/10/2017 and pre-processed as previously described^18^. Genes in the GTEx dataset containing fewer than 10 samples with a TPM > 1 were excluded from downstream analysis on the basis of unreliable quantification for very lowly expressed genes. We removed 4 outlier samples in the GTEx dataset that had sample-sample connectivity Z-scores of > 2 as previously described^58^. Our starting GTEx dataset consisted of 62 samples and 14577 genes.

RNA-sequencing data^13^ from postmortem cervical spinal cord (Brohawn et al.) was downloaded using the SRP accession number SRP064478 on 5/24/2019. RNA-sequencing data^22^ from laser capture microdissected (LCM) MNs primed for degeneration (Krach et al.) was downloaded using the SRP accession number SRP067645 on 6/27/2019. RNA-sequencing data^23^ from iPSC-derived motor neurons from ALS8 patients (Oliveira et al.) was downloaded using the SRP accession number SRP223674 on 10/18/2020. Reads were aligned to the reference genome GRCh37 using STAR (version 2.5.2b) and quantified using RSEM (version 1.3.0) with Gencode v19 annotations. Genes in the Brohawn et al., Krach et al., and Oliveira et al. datasets with zero variance in expression were excluded from further analysis. All expression data was quantified in TPM (transcript per million reads) and TPM values +1 were log_2_ transformed to stabilize their variance. The starting Brohawn et al., Krach et al., and Oliveira et al. datasets consisted of 33050, 37138, and 14171 genes respectively. Krach et al. metadata tables were transcribed from the manuscript and missing values for PMI (postmortem interval) were imputed as the mean of all available PMI values.

### Identification and regression of biological and technical confounders from gene expression datasets

Gene expression values were subjected to dimensionality reduction using principal component analysis (PCA) with data centering and scaling. The top four components for the GTEx dataset, which collectively captures 70.6% of the total variance, were assigned to be expression principal components (PCs) 1-4. The top five components for the Brohawn et al., Krach et al., and Oliveira et al. dataset, which collectively capture 55.9%, 42.3%, and 56.3% of the total variance respectively, were assigned to be expression principal components (PCs) 1-5. Sequencing Q/C metrics were calculated using PicardTools (version 2.5.0) and dimensionally reduced using PCA with centering and scaling. Sequencing Q/C metrics were aggregated manually for Brohawn et al. and Krach et al. datasets and aggregated using MultiQC^59^ for the Oliveira et al. dataset (**Supplementary Data 2-4**). The top five components of sequencing metrics for the GTEx, Brohawn et al., Krach et al., and Oliveira et al. datasets, which collectively capture 75.1%, 91%, 90.4%, and 89.0% of the total variance respectively, were selected and assigned to be sequencing principal components (seqPCs) 1-5.

Numerical covariates of technical or biological importance were correlated with the expression PCs using Spearman’s correlation. Binary covariates were numerically encoded and multi-level categorical covariates of technical or biological importance were binarized using dummy variables. The model fit (R^2^) was used to assess correlations. We identified technical or biological covariates with a meaningful influence on gene expression if they were correlated with an R > 0.3 to any of the top expression PCs.

For the GTEx expression dataset, a linear model was fit for each gene as follows: GTEX.Expr ∼ seqPC1 + seqPC2 + seqPC3 + seqPC4 + seqPC5 + RIN + DTHHRDY + AGE + SMCENTER + TRISCHD and the effect of all covariates were regressed out. RIN indicates RNA integrity number, DTHHRDY represents the Hardy Scale, SMCENTER represents the center for sampling, and TRISCHD represents ischemic time. Ethnicity was not included in the linear model because it was collinear with SMCENTER (**Supplementary Fig. S1c**).

For the Brohawn et al. dataset, a linear model was fit for each gene as follows: Expr ∼ seqPC1 + seqPC2 + seqPC3 + seqPC4 +seqPC5 + Ethnicity + Sex + Age + Disease and the effect of all covariates except disease were regressed out.

For the Krach et al. dataset, a linear model was fit for each gene as follows: Expr ∼ Diagnosis + Age + Gender + RIN + PMI + seqPC1 + seqPC2 + seqPC3 + seqPC4 + seqPC5 and the effect of all covariates except diagnosis were regressed out.

For the Oliveira et al. dataset, a linear model was fit for each gene as follows: Expr ∼ Disease + Age + Sex + Tissue + seqPC1 + seqPC2 + seqPC3 + seqPC4 + seqPC5 and the effect of all covariates except disease were regressed out.

### Weighted Gene Co-expression Network Analysis (WGCNA)

We used WGCNA^4,58^ to generate a gene co-expression network using the GTEx dataset. We chose a soft power of 10 using the function “pickSoftThreshold” in the WGCNA R package to achieve optimal scale free topology. The Topological Overlap Matrix (TOM) was generated using the function “TOMsimilarity” in the WGCNA R package. A hierarchical gene clustering dendrogram was constructed using the TOM-based dissimilarity and module selection was tested using multiple permutations of the tree cutting parameters: minimum module size, cut height, and deep split. Minimum module size (mms) was assessed at 50, 100, and 150. Module merging (dcor) was assessed at 0.1, 0.2, and 0.25. Deep split (DS) was assessed at 2 and 4. By visual inspection, we utilized the following parameters: minimum module size = 100, module merging = 0.1, and deep split = 2 to generate the network. We assigned unique module identifiers (SC.M1-13) for each module color except the grey module which represents genes that were not co-expressed.

Pairwise Spearman’s correlations were calculated using 1000 randomly sampled genes within each module and a matched background set. If a module contained less than 1000 genes, all genes in the module were sampled.

### Enrichment of gene sets in co-expression modules

We analyzed a collection of gene sets generated by previous genetic studies of ALS. For large effect size ALS risk genes, we obtained 50 ALS literature curated ALS risk genes from two studies^19,20^. For the determination of cell-type specificity, we obtained the top 50 differentially expressed genes per cell-type by fold change from single-nuclei RNA sequencing of postnatal day 2 mouse spinal cord^24^. For the human ALS spinal cord differentially expressed gene (DEG) enrichments, we obtained 265 DEGs (172 up- and 93 down-regulated) at FDR < 0.10 using the union of DESeq2 and EdgeR analyses^13^. For the dissected motor neurons primed for neurodegeneration, we obtained 1600 DEGs (1290 up- and 310 down-regulated) at FDR < 0.05 from the report^22^. For the iPSC-derived motor neurons from ALS patients, we chose to examine the severe ALS vs. control comparison and filtered gene biotype for protein-coding genes, obtaining 653 DEGs (364 up- and 289 down-regulated) at FDR < 0.01 from the report^23^. For DEGs from SOD1-G93A mice, we obtained genes with a Bayes Factor > 3 (corresponding to FDR = 0.1) identified through a spatial transcriptomics study^12^.

Logistic regression was performed to calculate an odds ratio and p-value for enrichment of each co-expression module with a test gene set. The p-values were FDR-corrected for the number of modules analyzed.

### Enrichment of common ALS genetic risk variants in co-expression modules

Stratified LD score regression^29^ was performed to evaluate the enrichment of common risk variants from GWAS studies of ALS^21^ (N=80,610), Inflammatory Bowel Disorder (IBD)^30^ (N=86,640), Autism Spectrum Disorder (ASD)^32,33^ (N=15,954), Alzheimer’s Disease (AD)^34^ (N=455,258), Frontotemporal Dementia (FTD)^35^ (N=12,928), and Progressive Supranuclear Palsy (PSP)^36^ (N=12,308) within each co-expression module. We used the full baseline model of 53 functional categories per calculation of partitioned heritability (1000 Genomes Phase 3). We defined genomic regions of each co-expression module as their constituent gene bodies plus a flanking region of +/- 10 KB.

### Gene Ontology Enrichments

Gene Ontology enrichment analysis was performed using the gProfiler package^60^ with hierarchical filtering set to “moderate”, domain size set to “annotated”, src filter set to “GO:BP” and “GO:MF”, electronic annotations (IEAs) included, maximum p-value set to 0.05, correction method set to “FDR”, and a maximum set size of 1000. To reflect the weight of each gene within a co-expression module, genes were ordered by their module membership (kME) during the enrichment calculations.

### Network visualization of module hub genes

For each module, an undirected and weighted network amongst the top 30 module hub genes (by kME) was visualized using the igraph package in R^61^.

### Protein-protein interaction (PPI) networks

PPI networks were produced using the Disease Association Protein-Protein Evaluator (DAPPLE)^62^. Two hundred and thirty eight of the top 300 SC.M4 hub genes, 238 of the top 300 SC.M2 hub genes, 84 of top 100 SC.M6 hub genes as ranked by Module Membership (kME) were found in the DAPPLE database and seeded into the shown PPI network (**Fig. 3e, 3f, 4b**). All networks were produced from 1,000 permutations (within-degree node-label permutation). SC.M6, SC.M4, and SC.M2 contain the following network properties respectively: direct edges count (p = 0.001, p =0.001, p = 0.001), seed direct degrees mean (p = 0.001, p = 0.001, p = 0.018), seed indirect degrees mean (p = 0.001, p = 0.006, p = 0.006), and the CI degrees mean (p = 0.001, p = 0.132, p = 0.332).

### Module preservation analyses

Module preservation for co-expression modules in the Brohawn et al. and Krach et al. datasets were performed as previously described using the modulePreservation function in the WGCNA R package^58^. We performed 200 permutation tests using a randomSeed of 1.

### Correlations of module eigengenes to biological and technical covariates

To evaluate the correlations of module eigengenes to biological and technical covariates in the Brohawn et al. and Krach et al. datasets, a Student p-value was calculated using the function “corPvalueStudent” in the WGCNA R package. P-values were FDR corrected across the modules and plotted using the function “labeledHeatmap” in the WGCNA R package. To account for multiple biological replicates per patient in the Oliveira et al. dataset, a mixed effect linear model was fitted for each module eigengene and each of the following covariates: Disease, Age, Sex, Tissue, seqPC1, seqPC2, seqPC3, seqPC4, and seqPC5. The covariate “Isolate”, which uniquely identifies each individual, was used as the random effect.

## Data Availability

Module assignments are provided as Supplementary Data 1 and the full range of values underlying the barplots and heatmaps in Figures 3a-b, 4a, 5a-c, and Supplementary Figures S4, S5d, S6d, S7d, S8, S9 are provided as a Source Data File.

## Code Availability

Underlying R code to run WGCNA is available at [https://github.com/dhglab/ALS-Gene-Network-Manuscript].

## Acknowledgements

We thank Christopher Hartl, William Pembroke, Timothy S. Chang, and other members of the Geschwind Laboratory for their technical assistance and manuscript feedback. We also thank Aaron D. Gitler and members of his laboratory for their stimulating discussions and assistance with the genetic modifier screen data. Funding for this work was provided by the Boyer Endowment Award, Helga K. & Walter Oppenheimer Endowment Award, and the MacDowell Endowment Award through the UCLA Undergraduate Research Center - Sciences (to J.C.W), NIMH 1F32MH114620 (to G.R), and NIMH 5R01MH109912 (to D.H.G).

## Author Contributions

J.C.W, G.R, and D.H.G planned the analyses and wrote the manuscript. Analyses were performed by J.C.W with supervision from G.R. and D.H.G.

