## Supplementary Material for "Gene co-expression network analysis in human spinal cord highlights mechanisms underlying amyotrophic lateral sclerosis susceptibility"

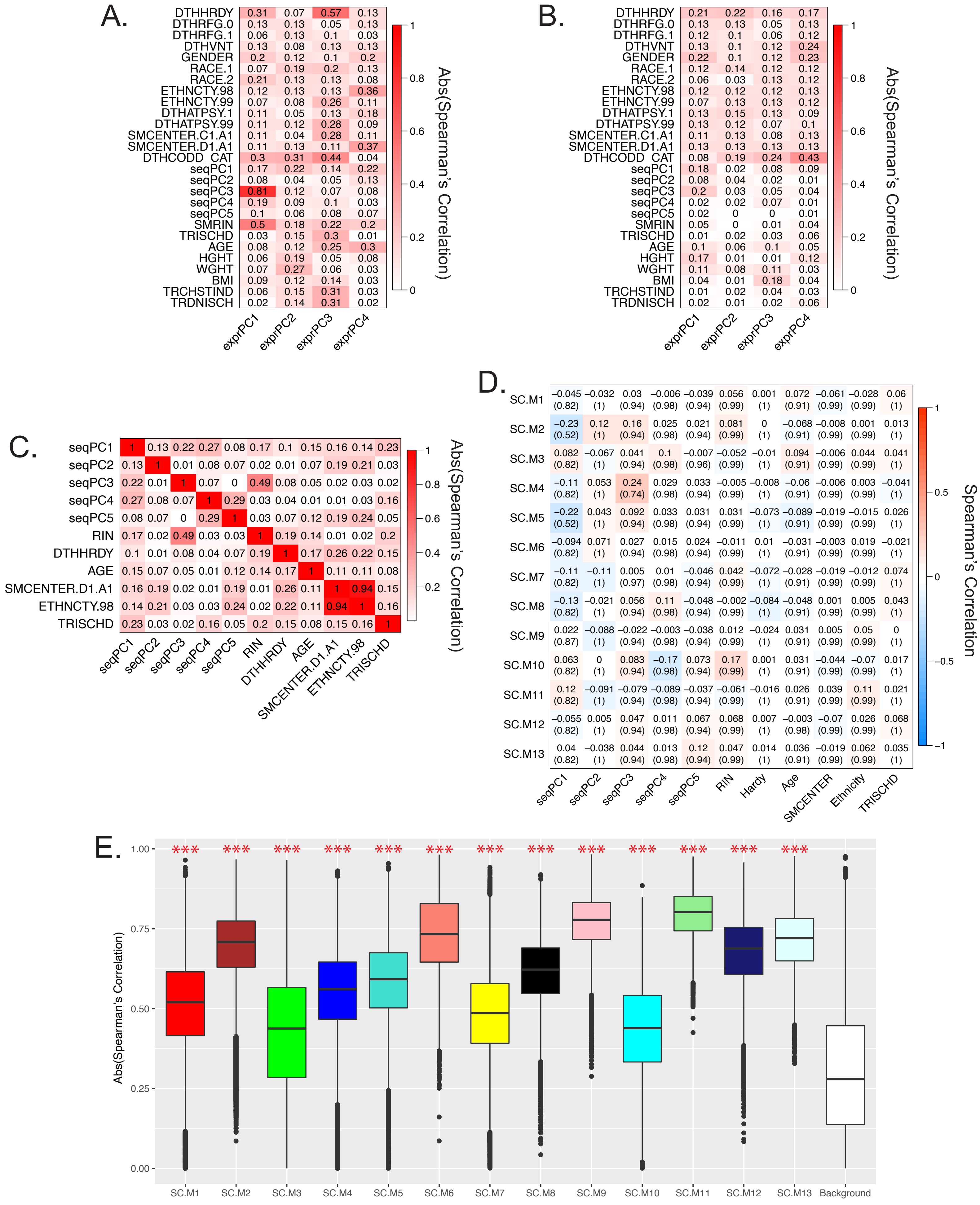
A.
DTHHRDY
DTHRFG.0
DTHRFG.1
DTHVNT
GENDER
RACE.1
RACE.2
ETHNCTY.98
ETHNCTY.99
DTHATPSY.1
DTHATPSY.99
SMCENTER.C1.A1
SMCENTER.D1.A1
DTHCODD_CAT
seqPC1
seqPC2
seqPC3
seqPC4
seqPC5
SMRIN
TRISCHD
AGE
HGHT
WGHT
BMI
TRCHSTIND
TRDNISCH
exprPC1
exprPC2
exprPC3
exprPC4
Abs(Spearman's Correlation)
B.
C.
seqPC1
seqPC2
seqPC3
seqPC4
seqPC5
RIN
DTHHRDY
AGE
SMCENTER.D1.A1
ETHNCTY.98
TRISCHD
D.
SC.M1
SC.M2
SC.M3
SC.M4
SC.M5
SC.M6
SC.M7
SC.M8
SC.M9
SC.M10
SC.M11
SC.M12
SC.M13
seqPC1
seqPC2
seqPC3
seqPC4
seqPC5
RIN
Hardy
Age
SMCENTER
Ethnicity
TRISCHD
Spearman's Correlation
E.
Abs(Spearman's Correlation)
SC.M1
SC.M2
SC.M3
SC.M4
SC.M5
SC.M6
SC.M7
SC.M8
SC.M9
SC.M10
SC.M11
SC.M12
SC.M13
Background

**Supplementary Figure S1.** Correction for technical and biological drivers of expression variance in the GTEX human spinal cord dataset.

(A) Spearman's correlations between the top four expression principal components (exprPCs1-4) and technical and biological covariates. RIN is RNA Integrity Number. DTHHRDY indicates Hardy's Death Classification which measures time from first onset of symptoms to death. TRISCHD indicates ischemic time as defined by the time of death to when tissue was fixed or frozen. Complete code annotations can be found in the GTEx Portal v7.

(B) Same as (A), except using the covariate-corrected human spinal cord expression matrix.

(C) Pairwise Spearman's correlations between biological and technical covariates in the human spinal cord dataset to check for covariate collinearity.

(D) Spearman's correlations of module eigengenes to traits in the covariate-corrected human spinal cord dataset. Significance was assessed using Student's t-test.

(E) Pairwise Spearman's correlations of 1,000 randomly sampled genes from each module and a background set. Significance of difference between each module with the background set was assessed using Student's t-test. *** (FDR corrected p-value < 0.001).

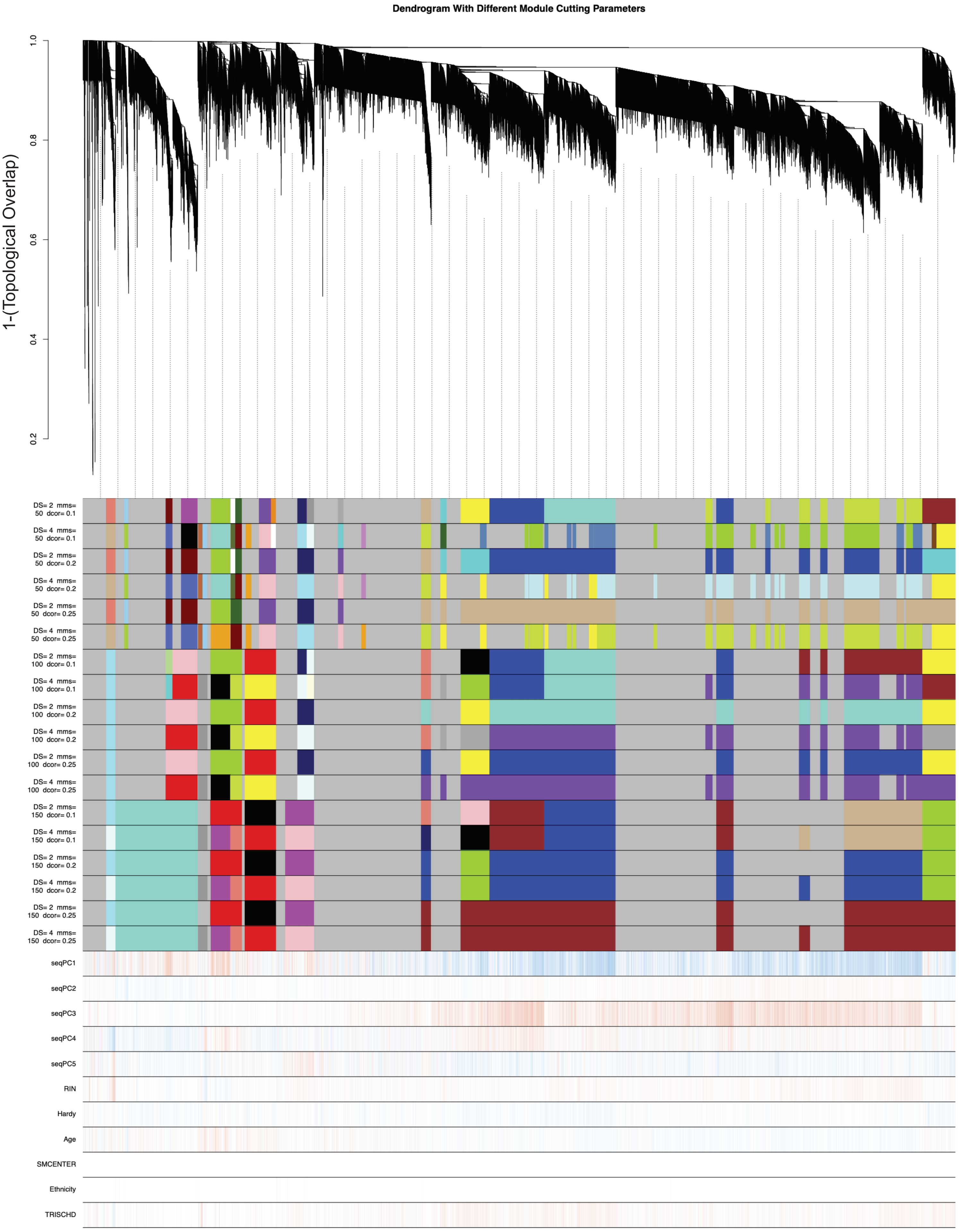

Dendrogram With Different Module Cutting Parameters
1-(Topological Overlap)
0.2
0.4
0.6
0.8
1.0
DS= 2 mms= 50 dcor= 0.1
DS= 4 mms= 50 dcor= 0.1
DS= 2 mms= 50 dcor= 0.2
DS= 4 mms= 50 dcor= 0.2
DS= 2 mms= 50 dcor= 0.25
DS= 4 mms= 50 dcor= 0.25
DS= 2 mms= 100 dcor= 0.1
DS= 4 mms= 100 dcor= 0.1
DS= 2 mms= 100 dcor= 0.2
DS= 4 mms= 100 dcor= 0.2
DS= 2 mms= 100 dcor= 0.25
DS= 4 mms= 100 dcor= 0.25
DS= 2 mms= 150 dcor= 0.1
DS= 4 mms= 150 dcor= 0.1
DS= 2 mms= 150 dcor= 0.2
DS= 4 mms= 150 dcor= 0.2
DS= 2 mms= 150 dcor= 0.25
DS= 4 mms= 150 dcor= 0.25
seqPC1
seqPC2
seqPC3
seqPC4
seqPC5
RIN
Hardy
Age
SMCENTER
Ethnicity
TRISCHD

**Supplementary Figure S2.** Optimization of tree cutting parameters to generate co-expression modules.

Dendrogram of the topological overlap matrix for gene co-expression dissimilarity. Co-expression module assignments are shown in each color track. Various permutations of minimum module size (mms), module merging (dcor), and deep split parameters (DS) were tested. Spearman correlations for each gene to biological and technical covariates are shown below the color tracks (Blue: Negative; Red: Positive).

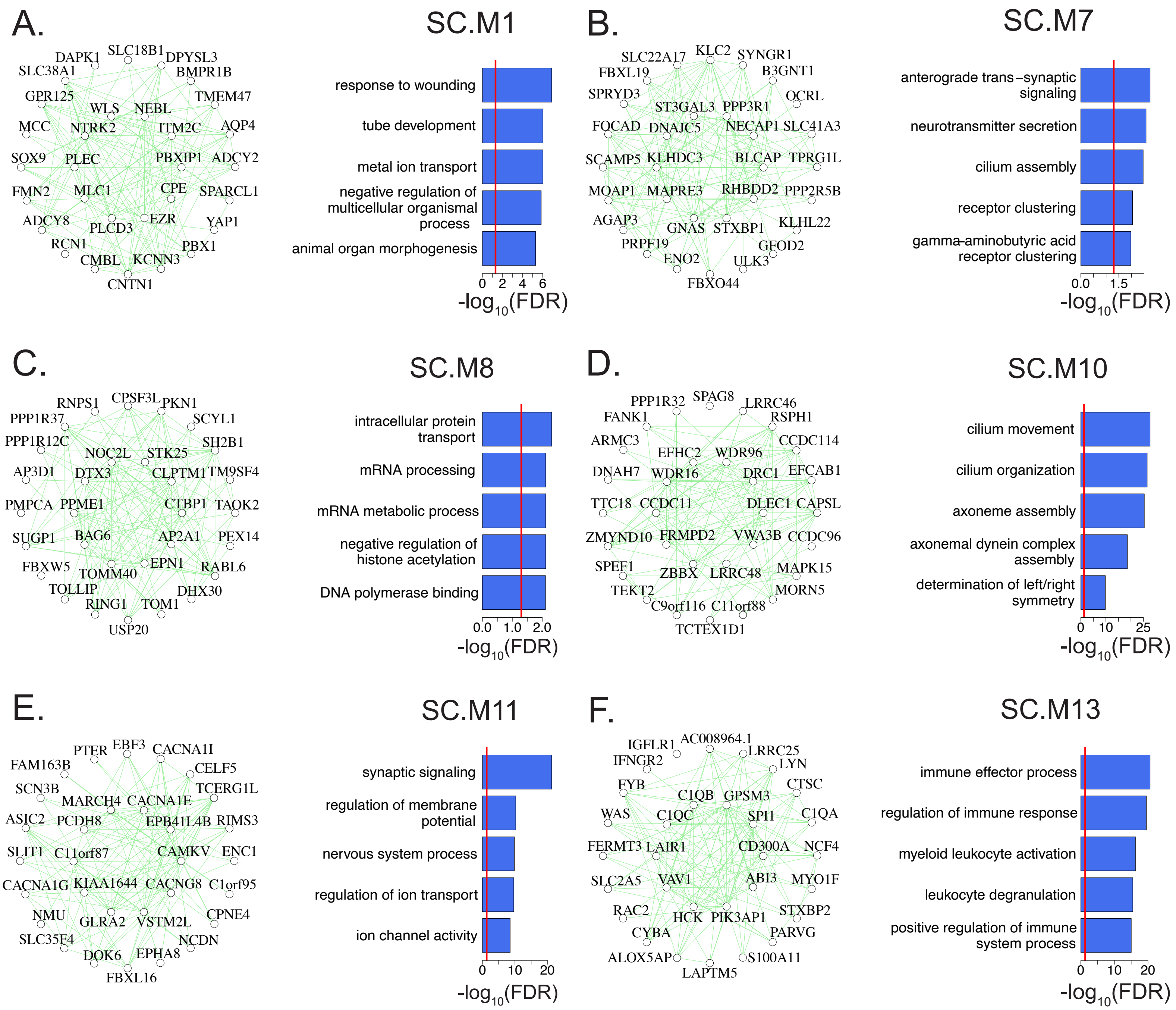

A.
SC.M1
DAPK1
SLC18B1
DPYSL3
SLC38A1
BMPR1B
GPR125
TMEM47
WLS
NEBL
MCC
NTRK2
ITM2C
AQP4
SOX9
PLEC
PBXIP1
ADCY2
FMN2
MLC1
CPE
SPARCL1
ADCY8
PLCD3
EZR
YAP1
RCN1
PBX1
CMBL
KCNN3
CNTN1
response to wounding
tube development
metal ion transport
negative regulation of multicellular organismal process
animal organ morphogenesis
0 2 4 6
-log10(FDR)
B.
SC.M7
SLC22A17
KLC2
SYNGR1
FBXL19
B3GNT1
SPRYD3
OCRL
ST3GAL3
PPP3R1
FOCAD
DNAJC5
NECAP1
SLC41A3
SCAMP5
KLHDC3
BLCAP
TPRG1L
MOAP1
MAPRE3
RHBDD2
PPP2R5B
AGAP3
GNAS
STXBP1
KLHL22
PRPF19
GFOD2
ENO2
ULK3
FBXO44
anterograde trans−synaptic signaling
neurotransmitter secretion
cilium assembly
receptor clustering
gamma-aminobutyric acid receptor clustering
0.0 1.5
-log10(FDR)
C.
SC.M8
RNPS1
CPSF3L
PKN1
PPP1R37
SCYL1
PPP1R12C
SH2B1
NOC2L
STK25
AP3D1
DTX3
CLPTM1
TM9SF4
PMPCA
PPME1
CTBP1
TAOK2
SUGP1
BAG6
AP2A1
PEX14
FBXW5
EPN1
RABL6
TOMM40
TOLLIP
DHX30
RING1
TOM1
USP20
intracellular protein transport
mRNA processing
mRNA metabolic process
negative regulation of histone acetylation
DNA polymerase binding
0.0 1.0 2.0
-log10(FDR)
D.
SC.M10
PPP1R32
SPAG8
LRRC46
FANK1
RSPH1
ARMC3
CCDC114
EFHC2
WDR96
DNAH7
WDR16
DRC1
EFCAB1
TTC18
CCDC11
DLEC1
CAPSL
ZMYND10
FRMPD2
VWA3B
CCDC96
SPEF1
ZBBX
LRRC48
MAPK15
TEKT2
MORN5
C9orf116
C11orf88
TCTEX1D1
cilium movement
cilium organization
axoneme assembly
axonemal dynein complex assembly
determination of left/right symmetry
0 10 25
-log10(FDR)
E.
SC.M11
PTER
EBF3
CACNA1I
FAM163B
CELF5
SCN3B
TCERG1L
MARCH4
CACNA1E
ASIC2
PCDH8
EPB41L4B
RIMS3
SLIT1
C11orf87
CAMKV
ENC1
CACNA1G
KIAA1644
CACNG8
C1orf95
NMU
GLRA2
VSTM2L
CPNE4
SLC35F4
NCDN
DOK6
EPHA8
FBXL16
synaptic signaling
regulation of membrane potential
nervous system process
regulation of ion transport
ion channel activity
0 10 20
-log10(FDR)
F.
SC.M13
IGFLR1
AC008964.1
LRRC25
IFNGR2
LYN
FYB
CTSC
C1QB
GPSM3
WAS
C1QC
SPI1
C1QA
FERMT3
LAIR1
CD300A
NCF4
SLC2A5
VAV1
ABI3
MYO1F
RAC2
STXBP2
HCK
PIK3AP1
CYBA
PARVG
ALOX5AP
S100A11
LAPTM5
immune effector process
regulation of immune response
myeloid leukocyte activation
leukocyte degranulation
positive regulation of immune system process
0 10 20
-log10(FDR)

**Supplementary Figure S3.** GO term annotations and top hubs of co-expression modules not displayed in the main figures.

(A, B, C, D, E, F) Top 30 hub genes and 300 connections for SC.M1, SC.M7, SC.M8, SC.M10, SC.M11, and SC.M13 respectively (left panels). Top 5 enriched GO terms for each of the respective modules (right panels). Red lines mark an FDR corrected p-value threshold of 0.05.

A.

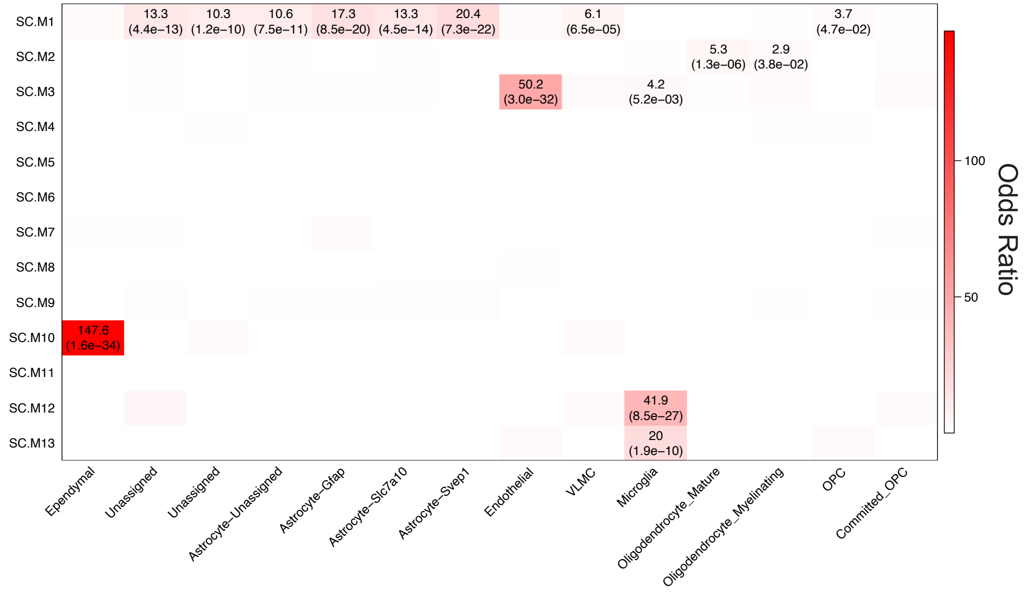
SC.M1
SC.M2
SC.M3
SC.M4
SC.M5
SC.M6
SC.M7
SC.M8
SC.M9
SC.M10
SC.M11
SC.M12
SC.M13
13.3 (4.4e−13)
10.3 (1.2e−10)
10.6 (7.5e−11)
17.3 (8.5e−20)
13.3 (4.5e−14)
20.4 (7.3e−22)
6.1 (6.5e−05)
3.7 (4.7e−02)
5.3 (1.3e−06)
2.9 (3.8e−02)
50.2 (3.0e−32)
4.2 (5.2e−03)
147.6 (1.6e−34)
41.9 (8.5e−27)
20 (1.9e−10)
Ependymal
Unassigned
Unassigned
Astrocyte−Unassigned
Astrocyte−Gfap
Astrocyte−Slc7a10
Astrocyte−Svep1
Endothelial
VLMC
Microglia
Oligodendrocyte_Mature
Oligodendrocyte_Myelinating
OPC
Committed_OPC
100
50
Odds Ratio

B.

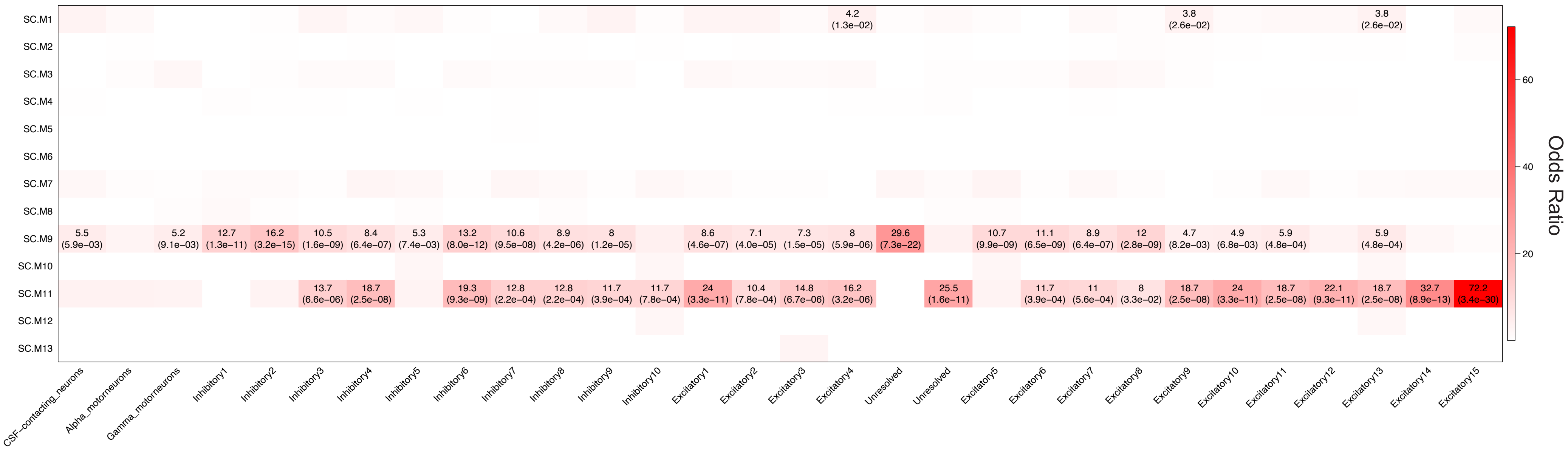
SC.M1
SC.M2
SC.M3
SC.M4
SC.M5
SC.M6
SC.M7
SC.M8
SC.M9
SC.M10
SC.M11
SC.M12
SC.M13
4.2 (1.3e−02)
3.8 (2.6e−02)
3.8 (2.6e−02)
5.5 (5.9e−03)
5.2 (9.1e−03)
12.7 (1.3e−11)
16.2 (3.2e−15)
10.5 (1.6e−09)
8.4 (6.4e−07)
5.3 (7.4e−03)
13.2 (8.0e−12)
10.6 (9.5e−08)
8.9 (4.2e−06)
8 (1.2e−05)
8.6 (4.6e−07)
7.1 (4.0e−05)
7.3 (1.5e−05)
8 (5.9e−06)
29.6 (7.3e−22)
10.7 (9.9e−09)
11.1 (6.5e−09)
8.9 (6.4e−07)
12 (2.8e−09)
4.7 (8.2e−03)
4.9 (6.8e−03)
5.9 (4.8e−04)
5.9 (4.8e−04)
13.7 (6.6e−06)
18.7 (2.5e−08)
19.3 (9.3e−09)
12.8 (2.2e−04)
12.8 (2.2e−04)
11.7 (3.9e−04)
11.7 (7.8e−04)
24 (3.3e−11)
10.4 (7.8e−04)
14.8 (6.7e−06)
16.2 (3.2e−06)
25.5 (1.6e−11)
11.7 (3.9e−04)
11 (5.6e−04)
8 (3.3e−02)
18.7 (2.5e−08)
24 (3.3e−11)
18.7 (2.5e−08)
22.1 (9.3e−11)
18.7 (2.5e−08)
32.7 (8.9e−13)
72.2 (3.4e−30)
CSF−contacting_neurons
Alpha_motorneurons
Gamma_motorneurons
Inhibitory1
Inhibitory2
Inhibitory3
Inhibitory4
Inhibitory5
Inhibitory6
Inhibitory7
Inhibitory8
Inhibitory9
Inhibitory10
Excitatory1
Excitatory2
Excitatory3
Excitatory4
Unresolved
Unresolved
Excitatory5
Excitatory6
Excitatory7
Excitatory8
Excitatory9
Excitatory10
Excitatory11
Excitatory12
Excitatory13
Excitatory14
Excitatory15
60
40
20
Odds Ratio

**Supplementary Figure S4.** Enrichment of cell-type markers from mouse spinal cord within co-expression modules.

(A) Enrichments of glial cell-type markers from mouse spinal cord[19] within co-expression modules. FDR corrected enrichment p-values < 0.05 are shown within the parentheses.

(B) Same as (A), except using neuronal cell-type markers.

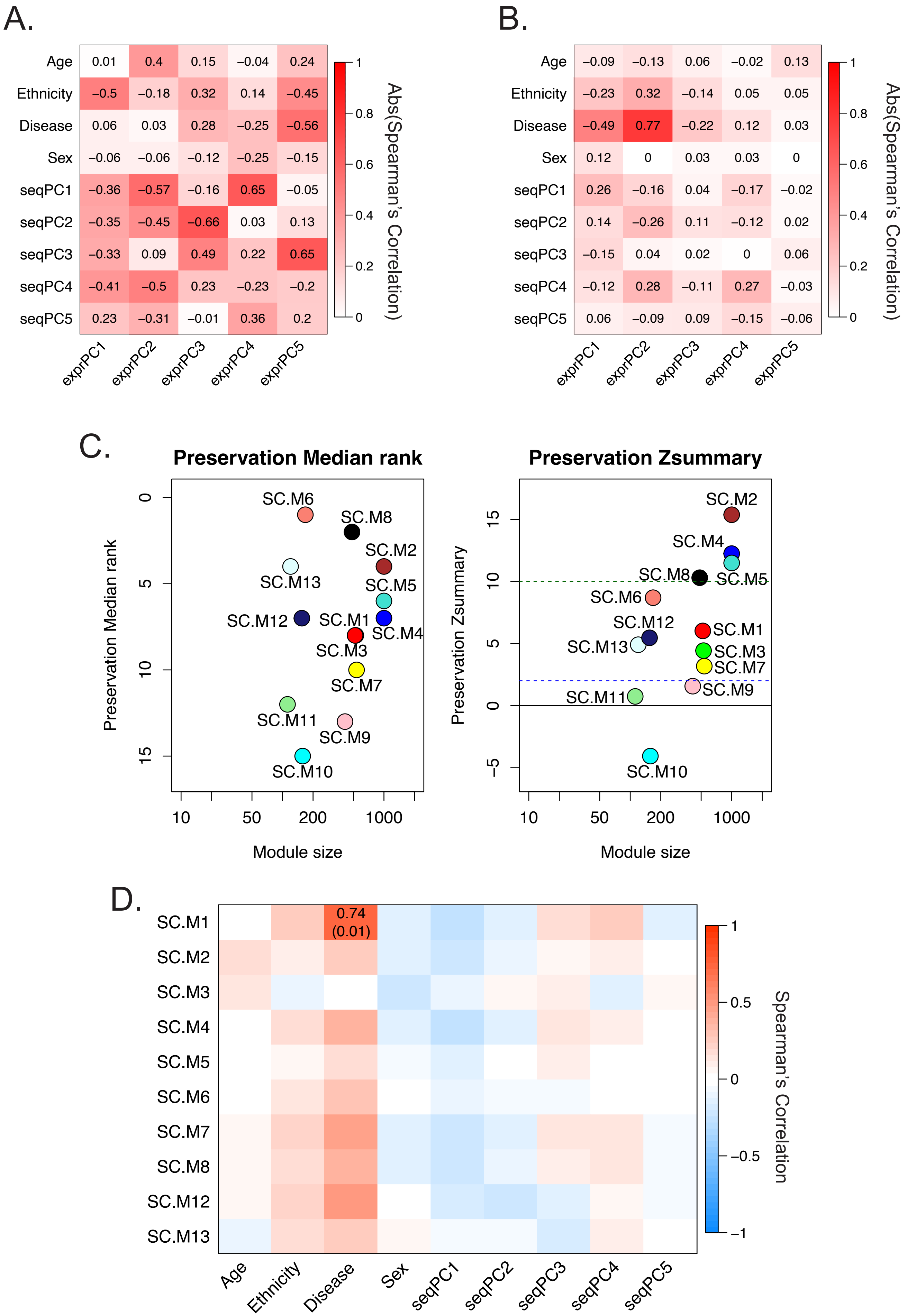
A.
Age 0.01 0.4 0.15 −0.04 0.24
Ethnicity −0.5 −0.18 0.32 0.14 −0.45
Disease 0.06 0.03 0.28 −0.25 −0.56
Sex −0.06 −0.06 −0.12 −0.25 −0.15
seqPC1 −0.36 −0.57 −0.16 0.65 −0.05
seqPC2 −0.35 −0.45 −0.66 0.03 0.13
seqPC3 −0.33 0.09 0.49 0.22 0.65
seqPC4 −0.41 −0.5 0.23 −0.23 −0.2
seqPC5 0.23 −0.31 −0.01 0.36 0.2
exprPC1 exprPC2 exprPC3 exprPC4 exprPC5
Abs(Spearman's Correlation)
B.
Age −0.09 −0.13 0.06 −0.02 0.13
Ethnicity −0.23 0.32 −0.14 0.05 0.05
Disease −0.49 0.77 −0.22 0.12 0.03
Sex 0.12 0 0.03 0.03 0
seqPC1 0.26 −0.16 0.04 −0.17 −0.02
seqPC2 0.14 −0.26 0.11 −0.12 0.02
seqPC3 −0.15 0.04 0.02 0 0.06
seqPC4 −0.12 0.28 −0.11 0.27 −0.03
seqPC5 0.06 −0.09 0.09 −0.15 −0.06
exprPC1 exprPC2 exprPC3 exprPC4 exprPC5
Abs(Spearman's Correlation)
C.
Preservation Median rank
Preservation Zsummary
Preservation Median rank
Preservation Zsummary
Module size
SC.M1 SC.M2 SC.M3 SC.M4 SC.M5 SC.M6 SC.M7 SC.M8 SC.M9 SC.M10 SC.M11 SC.M12 SC.M13
D.
SC.M1 SC.M2 SC.M3 SC.M4 SC.M5 SC.M6 SC.M7 SC.M8 SC.M12 SC.M13
0.74
(0.01)
Age Ethnicity Disease Sex seqPC1 seqPC2 seqPC3 seqPC4 seqPC5
Spearman's Correlation

**Supplementary Figure S5.** Covariate correction, module preservation, and module-trait correlations in postmortem human ALS spinal cord.

(A) Spearman's correlations between the top five expression principal components (exprPCs1-5) and technical and biological covariates.

(B) Same as (A), except using the covariate-corrected expression matrix.

(C) Module preservation analysis was used to calculate the $Z_{summary}$ statistic for each module[51] in postmortem human ALS spinal cord.

(D) Spearman's correlations of module eigengenes to traits correlated in postmortem human ALS spinal cord. Significance was assessed using Student's t-test. FDR corrected enrichment p-values < 0.05 are shown within the parentheses.

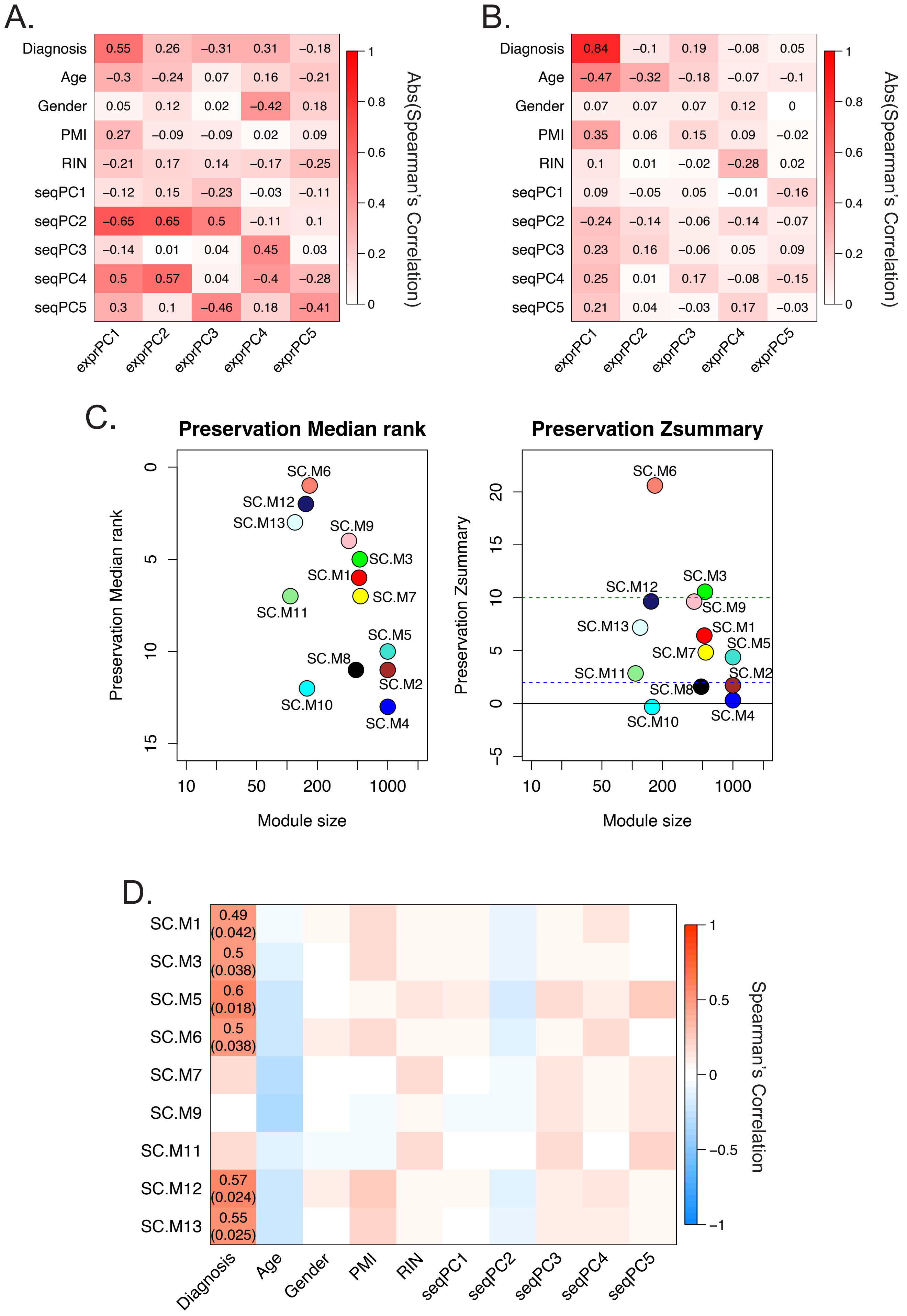
A.
B.
C.
Preservation Median rank
Preservation Zsummary
D.
Diagnosis
Age
Gender
PMI
RIN
seqPC1
seqPC2
seqPC3
seqPC4
seqPC5
exprPC1
exprPC2
exprPC3
exprPC4
exprPC5
Abs(Spearman's Correlation)
Module size
Spearman's Correlation
SC.M1
SC.M3
SC.M5
SC.M6
SC.M7
SC.M9
SC.M11
SC.M12
SC.M13
0.49 (0.042)
0.5 (0.038)
0.6 (0.018)
0.5 (0.038)
0.57 (0.024)
0.55 (0.025)

**Supplementary Figure S6**. Covariate correction, module preservation, and module trait correlations in presymptomatic motor neurons from human ALS spinal cord.

(A) Same as Supplementary Fig. S5a except using a dataset of presymptomatic motor neurons in the human ALS spinal cord. RIN is RNA Integrity Number. PMI is postmortem interval.

(B) Same as Supplementary Fig. S5b except using the covariate-corrected expression matrix generated from presymptomatic motor neurons in the human ALS spinal cord.

(C) Same as Supplementary Fig. S5c except module preservation was assessed in presymptomatic motor neurons from human ALS spinal cord dataset.

(D) Same as Supplementary Fig. S5d except using a dataset of presymptomatic motor neurons in the human ALS spinal cord. FDR corrected enrichment p-values < 0.05 are shown within the parentheses.

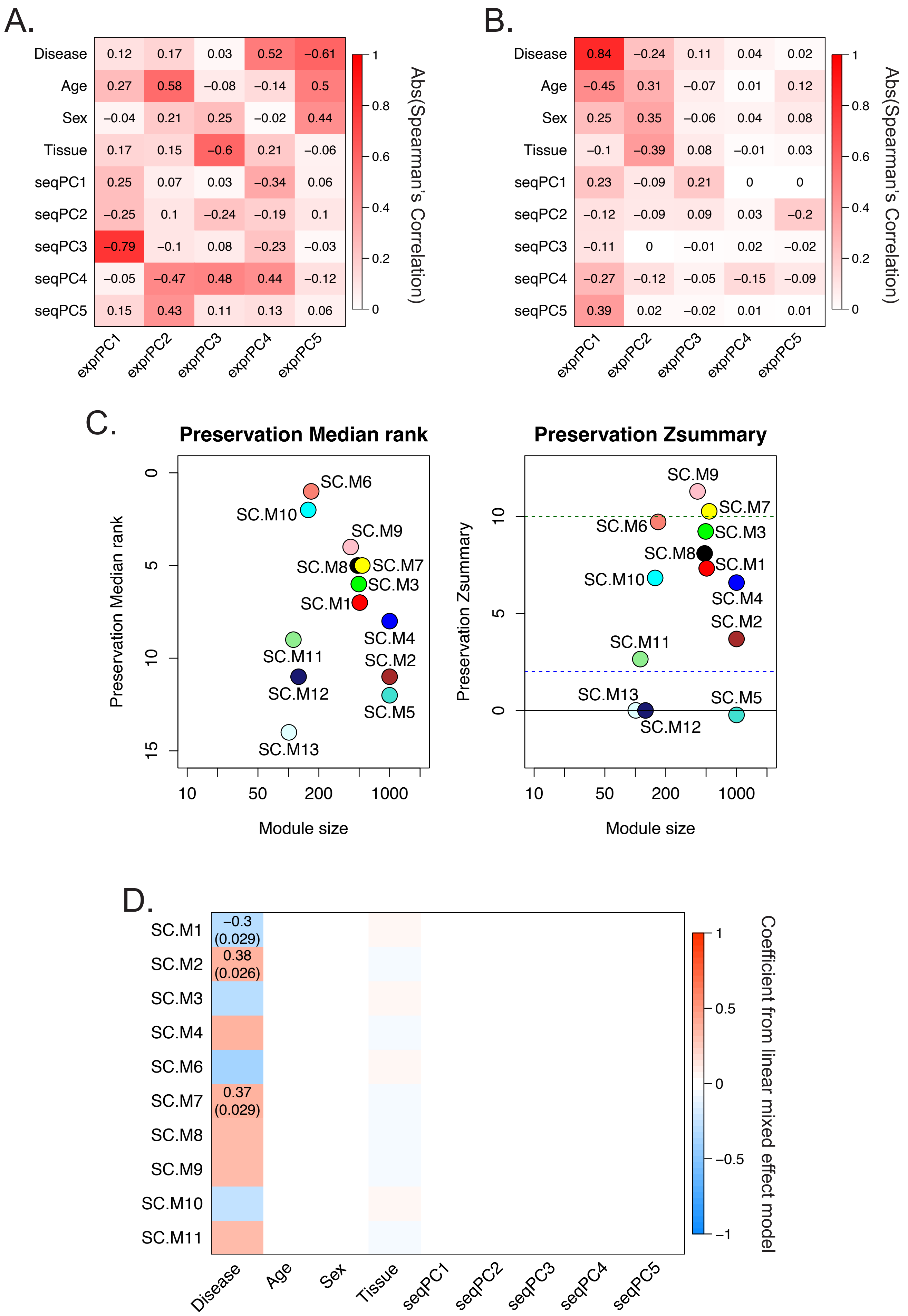
A.
Disease
Age
Sex
Tissue
seqPC1
seqPC2
seqPC3
seqPC4
seqPC5
exprPC1
exprPC2
exprPC3
exprPC4
exprPC5
Abs(Spearman's Correlation)
B.
C.
Preservation Median rank
Preservation Zsummary
Module size
D.
Coefficient from linear mixed effect model

**Supplementary Figure S7.** Covariate correction, module preservation, and module-trait correlations in iPSC-derived motor neurons from ALS8 patients.

(A) Same as Supplementary Fig. S5a except using a dataset of iPSC-derived motor neurons from ALS8 patients.

(B) Same as Supplementary Fig. S5b except using the covariate-corrected expression matrix generated from iPSC-derived motor neurons from ALS8 patients.

(C) Same as Supplementary Fig. S5c except module preservation was assessed in iPSC-derived motor neurons from ALS8 patients.

(D) Same as Supplementary Fig. S5d except using a dataset of iPSC-derived motor neurons from ALS8 patients. FDR corrected enrichment p-values $< 0.05$ are shown within the parentheses.

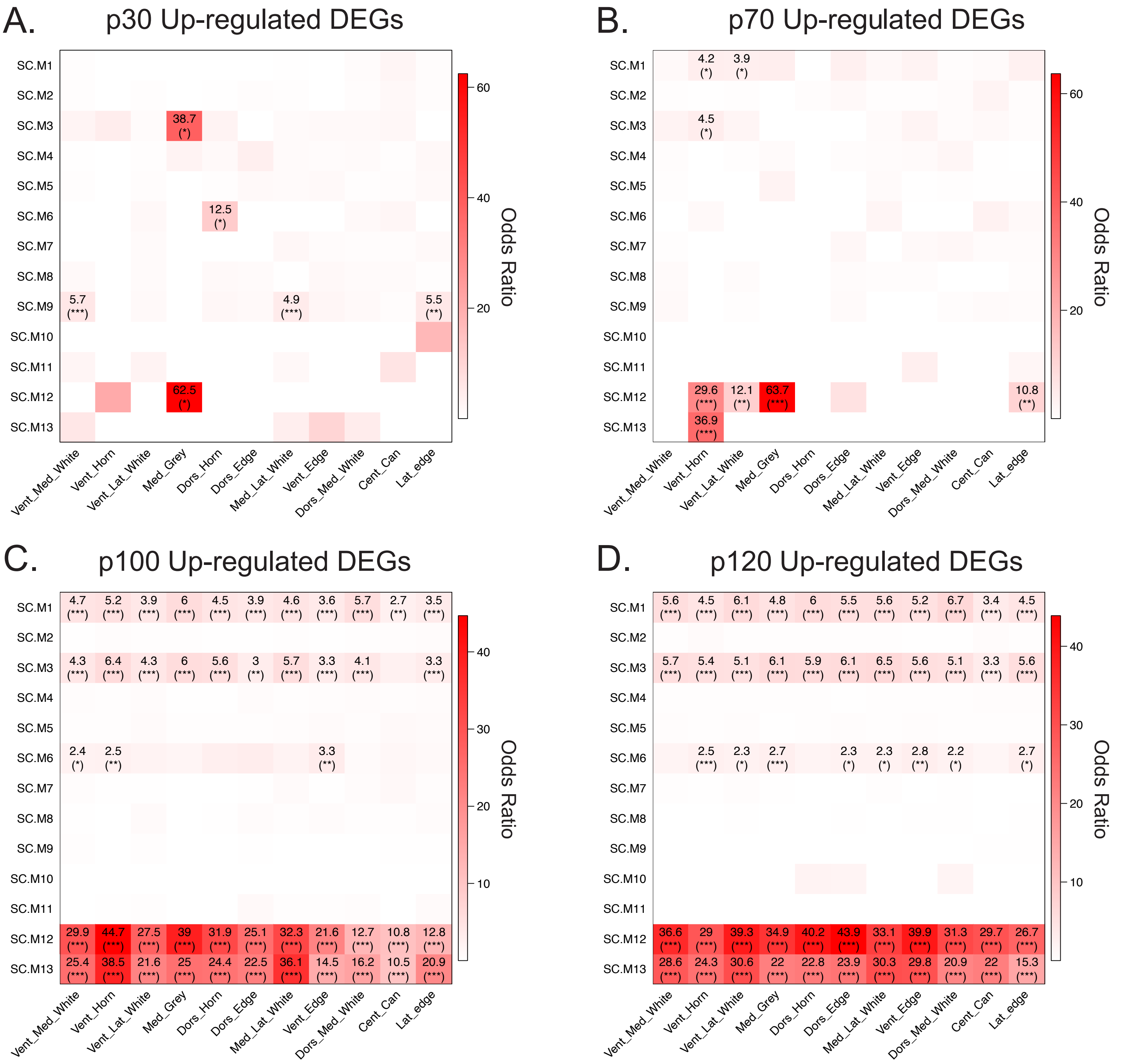
A.
p30 Up-regulated DEGs
B.
p70 Up-regulated DEGs
C.
p100 Up-regulated DEGs
D.
p120 Up-regulated DEGs
Odds Ratio
SC.M1
SC.M2
SC.M3
SC.M4
SC.M5
SC.M6
SC.M7
SC.M8
SC.M9
SC.M10
SC.M11
SC.M12
SC.M13
Vent_Med_White
Vent_Horn
Vent_Lat_White
Med_Grey
Dors_Horn
Dors_Edge
Med_Lat_White
Vent_Edge
Dors_Med_White
Cent_Can
Lat_edge

**Supplementary Figure S8.** Enrichment of ALS up-regulated genes for each Anatomical Annotated Region (AAR) from spatial transcriptomics study using SOD1-G93A mice in co-expression modules.

(A) Enrichment of up-regulated DEGs at stage p30 within co-expression modules. Only enrichments with an FDR corrected p-value < 0.05 are displayed. * (FDR corrected p-value < 0.05). ** (FDR corrected p-value < 0.01). *** (FDR corrected p-value < 0.001).

(B, C, E) Same as (A) except for the p70, p100, and p120 mouse respectively.

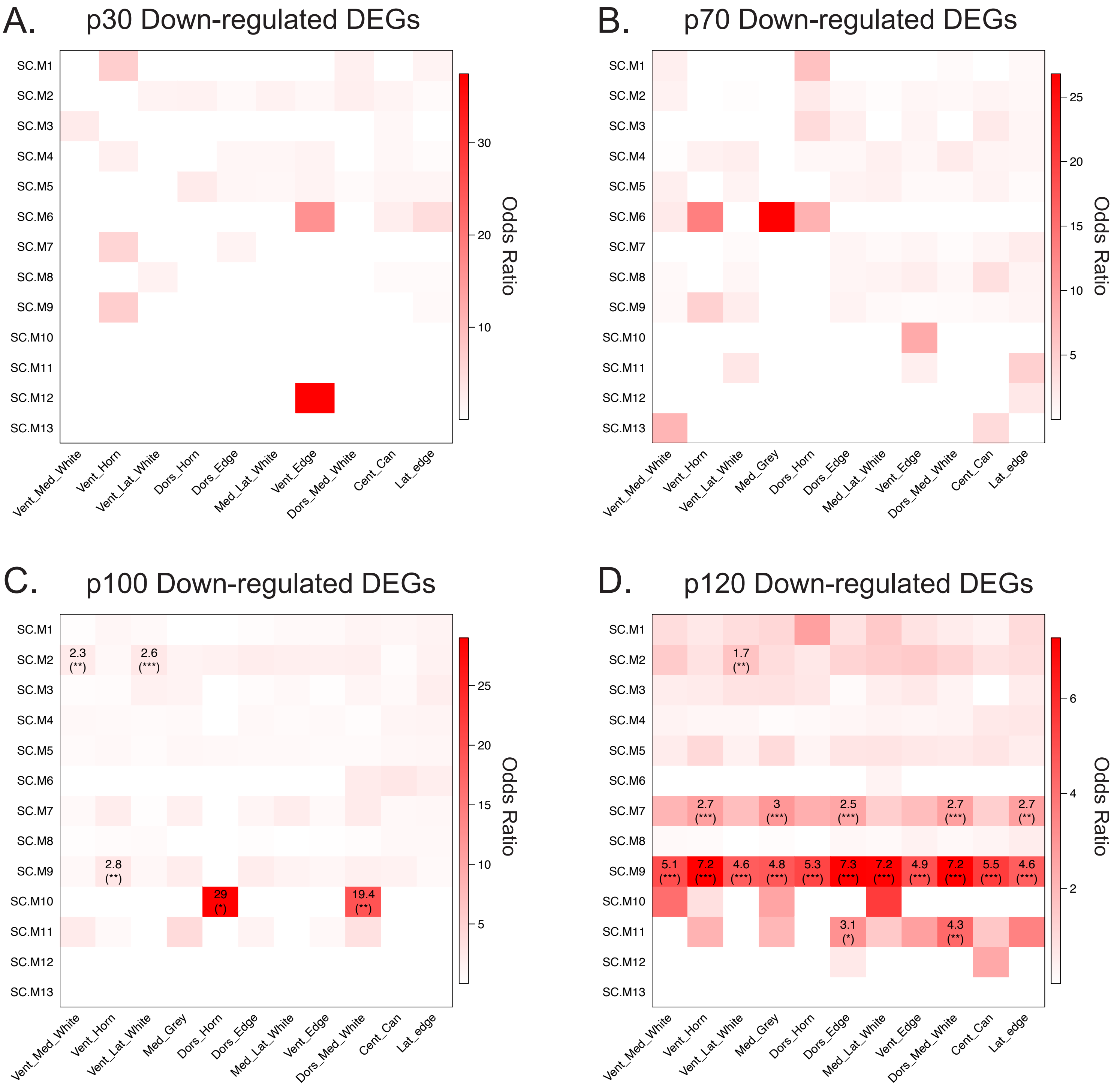
A.
p30 Down-regulated DEGs
B.
p70 Down-regulated DEGs
C.
p100 Down-regulated DEGs
D.
p120 Down-regulated DEGs
SC.M1
SC.M2
SC.M3
SC.M4
SC.M5
SC.M6
SC.M7
SC.M8
SC.M9
SC.M10
SC.M11
SC.M12
SC.M13
Vent_Med_White
Vent_Horn
Vent_Lat_White
Med_Grey
Dors_Horn
Dors_Edge
Med_Lat_White
Vent_Edge
Dors_Med_White
Cent_Can
Lat_edge
Odds Ratio
2.3 (**)
2.6 (***)
2.8 (**)
29 (*)
19.4 (**)
1.7 (**)
2.7 (***)
3 (***)
2.5 (***)
2.7 (***)
2.7 (**)
5.1 (***)
7.2 (***)
4.6 (***)
4.8 (***)
5.3 (***)
7.3 (***)
7.2 (***)
4.9 (***)
7.2 (***)
5.5 (***)
4.6 (***)
3.1 (*)
4.3 (**)

**Supplementary Figure S9.** Enrichment of ALS down-regulated genes for each Anatomical Annotated Region (AAR) from spatial transcriptomics study using SOD1-G93A mice in co-expression modules.

(A) Enrichment of down-regulated DEGs at stage p30 within co-expression modules. Only enrichments with an FDR corrected p-value < 0.05 are displayed. * (FDR corrected p-value < 0.05). ** (FDR corrected p-value < 0.01). *** (FDR corrected p-value < 0.001).

(B, C, E) Same as (A) except for the p70, p100, and p120 mouse respectively.
